# A phage nucleus-associated protein from the jumbophage Churi inhibits bacterial growth through protein translation interference

**DOI:** 10.1101/2024.06.15.599175

**Authors:** Wichanan Wannasrichan, Sucheewin Krobthong, Chase J Morgan, Emily G Armbruster, Milan Gerovac, Yodying Yingchutrakul, Patompon Wongtrakoongate, Jörg Vogel, Chanat Aonbangkhen, Poochit Nonejuie, Joe Pogliano, Vorrapon Chaikeeratisak

## Abstract

Antibacterial proteins inhibiting *Pseudomonas aeruginosa* have been identified in various phages and explored as antibiotic alternatives. Here, we isolated a phiKZ-like phage, Churi, which encodes 364 open reading frames. We examined 15 early-expressed phage proteins for their ability to inhibit bacterial growth, and found that gp335, closely related to phiKZ-gp14, exhibits antibacterial activity. Similar to phiKZ-gp14, recently shown to form a complex with the *P. aeruginosa* ribosome, we predict experimentally that gp335 interacts with ribosomal proteins, suggesting its involvement in protein translation. GFP-tagged gp335 clusters around the phage nucleus as early as 15 minutes post-infection and remains associated with it throughout the infection, suggesting its role in protein expression in the cell cytoplasm. CRISPR-Cas13-mediated deletion of gp355 reveals that the mutant phage has a prolonged latent period. Altogether, we demonstrate that gp335 is an antibacterial protein of nucleus-forming phages that associates with the ribosomes at the phage nucleus.

## Introduction

*Pseudomonas aeruginosa* is a gram-negative opportunistic bacterium responsible for nosocomial infection and a leading cause of morbidity and mortality in immunocompromised and elderly patients^1–4^. The treatment of *P. aeruginosa* infection tends to be less successful compared to other bacterial infections due to high levels of both intrinsic and acquired antibiotic resistance^5,6^. Unfortunately, the slow rate of novel drug discovery and the dramatic evolution of antibiotic resistance increases the challenge of managing *P. aeruginosa* infections. Bacteriophages are increasingly being focused on as a promising alternative for dealing with the antimicrobial resistance crisis^7^. One concern with natural phage therapy is the possibility that genes for toxin production or antibiotic resistance can be accidentally packaged into phage head and transmitted to another bacterial host^8^. Therefore, alternatives to whole phage therapy may be preferred. Fortunately, application of phages is not only limited to the use of replicative phages, but also extendable to phage-derived products that can target host metabolic machinery and induce host cell lysis^9^.

Phages rely on many host factors to propagate, including proteins involved in DNA replication, RNA transcription, protein translation, and energy metabolism. Phages often encode proteins that hijack these mechanisms and optimize the intracellular environment for phage progeny production^10^. These proteins often target essential host proteins involved in gene expression and/or cell division^10^. Several of these proteins also show antibacterial activity^9,11^. However, phage proteins that hinder host protein translation are rarely found^10,12,13^. A number of studies have found that expressing phage-encoded proteins diminished bacterial growth after induction^14–20^. For example, *Pseudomonas* phage LUZ24 produces a small polypeptide called Igy, that specifically binds to the host DNA gyrase. The Igy protein exhibits antimicrobial activity against *P. aeruginosa* in both PAO1 and PA14 strains when expressed inside bacterial cells because it interrupts host DNA gyrase activity leading to the collapse of DNA replication and cell death^17,18^. ORF104 from *S. aureus* phage 77 also inhibits bacterial growth by binding host DnaI (helicase loader)^14^. Furthermore, some phages have been found to interfere with host RNA transcription. Phage LUZ19 encodes a protein, gp25.1 that interacts with β′-subunit of *P. aeruginosa* RNA polymerase. This protein inhibits host transcription and decreases bacterial growth^15,20^. Another example is gp12 encoded by phage 14-1, which inhibits transcription from host cell promoters by interacting with the α-subunit of *P. aeruginosa* RNA polymerase and may alter promoter specificity toward phage promoters^15^. These proteins however come from small genome phages. The ability of nucleus-forming jumbophage proteins to suppress bacterial growth remains largely unexplored.

Here, we identify Churi as a previously uncharacterized nucleus-forming jumbophage that is closely related to phiKZ and discover that it encodes an early gene product with antibacterial activity. Growth inhibition assay of Churi early expressed proteins shows that gp335 strongly inhibits the growth of *P. aeruginosa*, when overexpressed from a plasmid. Co-immunoprecipitation experiments suggest that gp335 interacts with ribosomes, which provides a plausible mechanism of action for this growth inhibition. During Churi infection, gp335 accumulates around the phage nucleus and remains associated as a cloud surrounding the nucleus throughout the infection. While a gp335-Churi loss of function mutant exhibits delay in bacterial cell lysis, complementation of the wildtype gp335-Churi can improve the lysis time. This study of the nucleus-forming jumbophage Churi reveals an early gene product that both suppresses bacterial growth likely through protein translation interference and plays a role in the phage replication cycle.

## Results

### Characteristics of phage Churi reveal that it is a nucleus-forming jumbophage

To derive a bacteriophage as a potential source of antibacterial proteins, a soil sample was collected from the area of Chulalongkorn University and incubated with *P. aeruginosa* PAO1. The supernatant of this culture was plated on *P. aeruginosa* and small plaques with morphology consistent with a jumbophage were purified. Churi plaque morphology is clear throughout the plaque area with rough borders and a plaque size is around 1-1.5 mm in diameter on 0.35% LB agar **(Figure 1a)**. Negative stain electron microscopy shows that the length of a Churi particle from the top head to the bottom of base plate is ∼300 nm while head alone is ∼120 nm in length and width. The tail is ∼140 nm in length and 25 nm in width **(Figure 1b)**. The latent period of phage Churi is ∼65 minutes post infection (mpi) at which point the number of phage progenies steeply rises. The average burst size of phage Churi is around 59 particles per cell **(Figure 1c)**. Fluorescence microscopy of infected cells shows a bright punctum of DAPI-stained phage DNA located at the cell pole around 10 mpi. The DNA punctum grows larger and migrates toward the center of the infected cell by 45 mpi **(Figure 1d and Supplementary Figure 1)**. During late infection, DAPI-stained horseshoe-shaped structures that were composed of mature virions, also called phage bouquets^21^, are found at the sides of central DNA at 90 mpi and becomes larger and brighter at 120 mpi. The number of infected cells increased throughout the infection time from 70% at 90 mpi to around 90% at 120 mpi **(Figure 1d and Supplementary Figure 1)**. This frequency of bouquet formation in Churi is highly similar to what is found in phage phiPA3 (∼80%) and is substantially prominent compared to phage phiKZ (∼23%) and 201phi2-1 (∼2%), suggesting the different degree of subcellular organization for phage maturation among them^21^.

**Figure 1.**
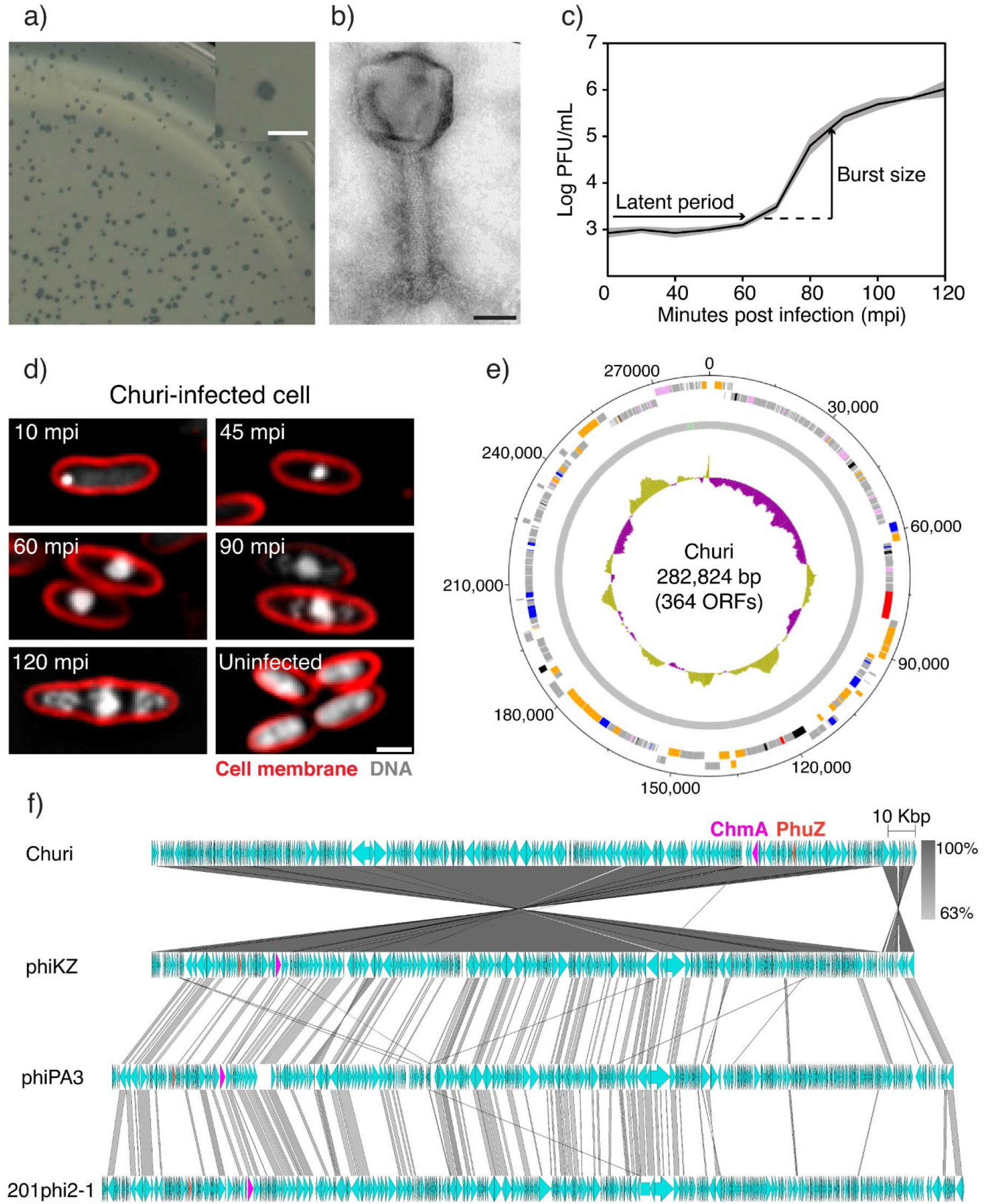
Characteristics of phage Churi reveal that it is a nucleus-forming jumbo phage. Characteristics of phage Churi. **a)** Double layer agar shows plaques morphology of Churi, scale bar represents 2 mm; **b)** A transmission electrogram reveals Churi particle morphology, which is categorized as a myovirus, and scale bar represents 50 mm. **c)** One-step growth curve of Churi. **d)** Single cell infection against *P. aeruginosa* of Churi under fluorescence microscope. The red of FM4-64 represents bacterial cell membrane while gray of DAPI represents DNA. A scale bar represents 1 µm. **e)** Circular genomic map of phage Churi. ORFs indicated by colors as follows: blue, DNA replication, transcription and translation and DNA repair; pink, nucleotide metabolism and modification; orange, virion structural and assembly; red, host-phage interaction and host lysis protein; black, others; and gray, hypothetical proteins. An inner plot presents the GC content across the genome (yellow above average and purple below average). The numbers surrounding the map represent numbers of nucleotides. The map was generated using Artemis: DNAPlotter version 18.1.0. **f)** Comparative genomic analysis shows the gene organization between phage Churi, phiKZ, phiPA3, and 201phi2-1 respectively. Red labelled arrows represent the location and orientation of PhuZ while magenta arrows represent shell (ChmA).

The complete genome of phage Churi was obtained by whole genome sequencing. The genome topology of Churi is double-stranded DNA, which is circularly permutated and terminally redundant. Genome size is 282,824 base pairs with a low GC content at 36.9% and encodes 364 open reading frames (ORFs) **(Figure 1e)**. Churi classifies as a jumbophage, which is defined to have a genome larger than 200 kb^22^. Churi encodes 7 tRNAs responsible for 6 amino acids: Threonine, Leucine, Proline, Isoleucine, Aspartic acid, and Asparagine. There are 282 ORFs that encode for hypothetical proteins with no known function, while 82 ORFs can be functionally annotated. Genes with known function are involved in the following processes: virion structural and assembly (42 genes); DNA replication, transcription and translation, and DNA repair (14 genes); nucleotide metabolism and modification (14 genes); host-phage interaction and host lysis protein (2 genes); and others (10 genes). Importantly, the nuclear shell protein ChmA and tubulin PhuZ are also found in Churi genome **(Figure 1f; pink and orange arrows)**. This indicates a common genotype of a bacteriophage in the family *Chimalliviridae,* which includes all known nucleus-forming phages^23^. VIRIDIC analysis^24^ between Churi and well-studied jumbophages (phiKZ, OMKO1, phiPA3, and 201phi2-1), that form nucleus-like structures shows that Churi is closely related to phiKZ and OMKO1 (another phiKZ-like phage) with 94.2% intergenomic sequence similarity **(Supplementary Table 1).** However, they are not the same species according to the demarcation criteria of ICTV^24,25^. In contrast, the sequence identity of Churi against phiPA3 and 201phi2-1 is 18.3% and 12.3% respectively. The orientation of most ORFs between Churi and phiKZ genomes is in opposite direction with some switching loci even though their genomic organization is highly similar. Lower numbers of Churi genes were found to match with phiPA3 and 201phi2-1 **(Figure 1f)**.

### gp335 is an early-expressed Churi protein that inhibits *P. aeruginosa* growth and interacts with host translation machinery

To test our hypothesis if nucleus-forming phage encode early genes with antimicrobial properties, we performed mass spectrometry on early infection cultures to identify candidates. We harvested infected cells from 2 infection methods: plate cultures and liquid cultures, at 15 mpi. While 27 non-virion phage proteins were identified during plate infection and 112 non-virion phage proteins were identified during liquid infection **(Supplementary Table 2 and 3)**, only 15 proteins were found in both experiments, assuring the presence of these phage-expressed proteins regardless of infection models. Even though these 15 proteins are annotated as hypothetical proteins, gp270 which is the homolog of gp068-phiKZ, a part of non-virion RNA polymerase complex (nvRNAP), are also detected. The nvRNAP is known as the product of early phage genes^26,27^ **(Table 1)**, supporting our proteomic results that show early proteins expressed from Churi. We then tested these 15 candidate proteins for growth inhibition against *P. aeruginosa* PAO1 strain by inducing phage gene expression on pHERD30T plasmids with arabinose. Empty-pHERD30T was used as a negative control while gp10 of phage JJ01^28^, whose homolog was previously identified as a strong bacterial growth inhibitor^15^, was used as a positive control. Of our candidates, only gp335 reduced the growth of *P. aeruginosa* when its expression was induced with arabinose for 18 hours. The reduction in colony forming units (CFUs) was at least 6 log fold compared to the non-induction culture **(Figure 2a and 2b)**. Homologs of gp335 in other *Pseudomonas* jumbophages (gp014-phiKZ and gp122-phiPA3) were also tested for the activity. Interestingly, only gp014-phiKZ exhibits inhibitory activity against *P. aeruginosa* PAO1 similar to gp335-Churi but gp122-phiPA3 has no growth inhibitory activity against the host **(Figure 2b)**. The realtime effect of gp335-Churi against *P. aeruginosa* was also observed by optical density (OD) in liquid culture at various levels of induction. During 10 hours of growth, 0.2% arabinose induction slightly lowered the optical density, 0.4% arabinose induction greatly lowered optical density, and 0.8% arabinose induction almost completely abrogated culture growth **(Figure 2c).** Likewise, this result pattern can be observed in the cells expressing gp014-phiKZ **(Figure 2d)**, suggesting the conserved antimicrobial activity between the homologs in Churi and phiKZ. The bacterial CFU is also found to sequentially decrease when inducing gp335-Churi at increasing concentration of arabinose **(Supplementary Figure 2)**, suggesting that the growth inhibition activity of gp335-Churi against *P. aeruginosa* is dose dependent. In contrast, there is no difference in the bacterial growth between uninduced and induced conditions for gp122-phiPA3 **(Figure 2e)**. The growth of empty-pHERD30T (negative control) containing bacteria does not decrease when induced with highest concentration of arabinose while gp10-JJ01 (positive control) does suppress growth even induced at the lowest arabinose concentration, indicating that the expression system functions as expected, and that growth suppression is truly specific to the expressed protein in cells **(Supplementary Figure 2)**.

**Figure 2.**
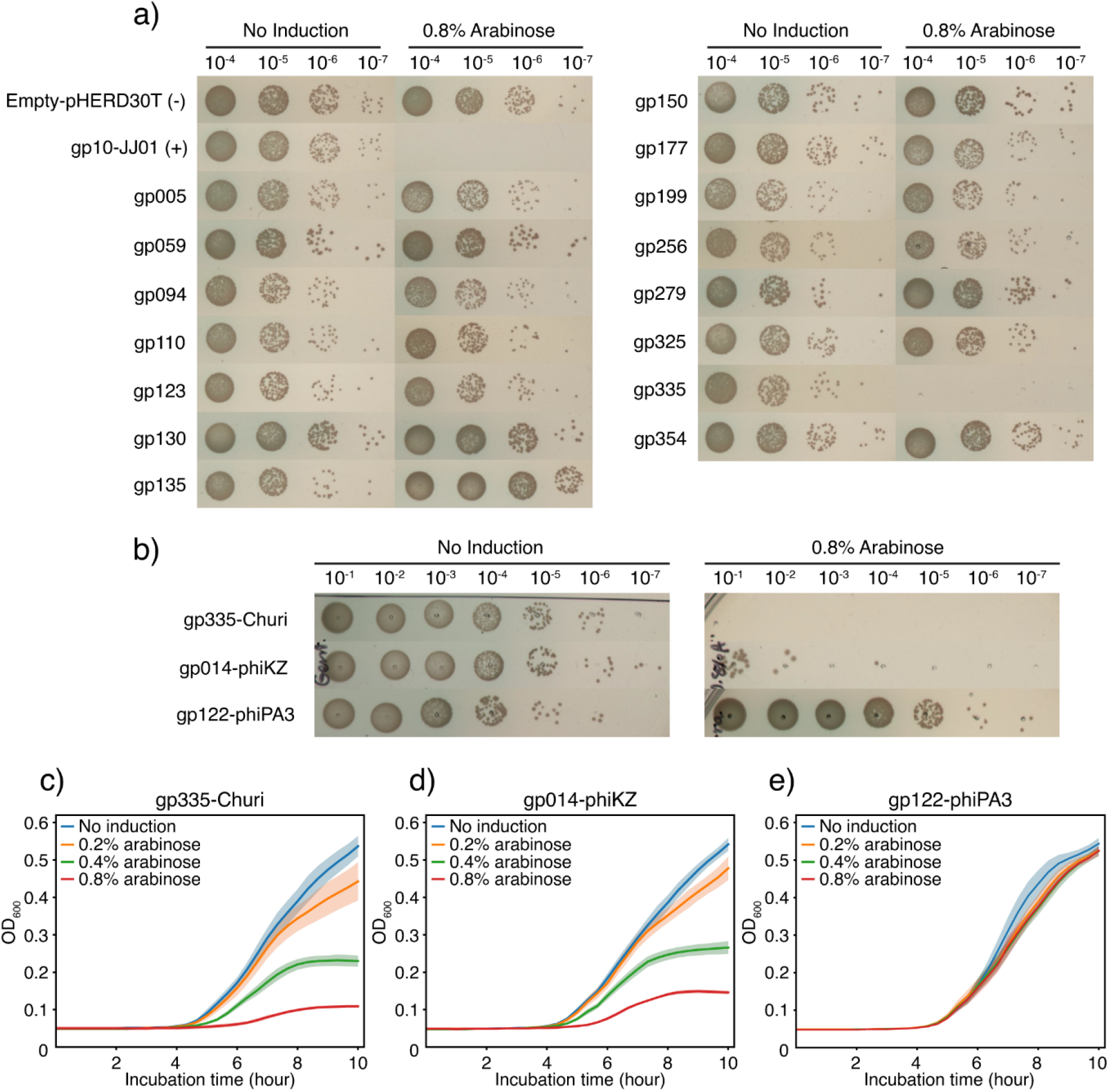
Growth inhibition assay of Churi-gene expressing *P. aeruginosa* reveals that gp335 substantially suppresses the bacterial growth. **a)** Churi genes that have inhibitory activity against *P. aeruginosa* PAO1 were screened from 15 proteins found to express in the early infection of both proteomic experiments. Empty-pHERD30T represents negative control while gp10-phage JJ01 is used as the positive control of growth inhibitory gene. Ten-fold dilution of gene-expressing bacterial culture is spotted on LB agar supplemented with gentamicin. The spots are shown as the dilution from 10^-4^ to 10^-7^. **b)** Inhibitory activity of gp335-Churi homologs in phiKZ and phiPA3. The spots are shown as the dilution from 10^-1^ to 10^-7^. All recombinant strains of *P. aeruginosa* are induced with or without 0.8% arabinose. **c)-e)** Optical density (OD_600_) of bacteria expressing gp335-Churi compared to gp014-phiKZ, and gp122-phiPA3 when the cells were induced with different concentration of arabinose (0.2, 0.4, and 0.8%). The OD_600_ values were measured every 20 minutes for 10 hours of incubation. Shaded error bar represents standard deviation (±SD) of n=6. **c)** gp335-Churi, **d)** gp014-phiKZ, and **e)** gp122-phiPA3.

**Table 1.** Selected non-structural proteins of Churi found to express at 15 minutes post infection from both proteomic experiments and their homologs in other nucleus-forming phages (phiKZ and phiPA3).

| Proteins | Function | Homologs |  |
| --- | --- | --- | --- |
|  |  | phiKZ | phiPA3 |
| gp005 | Hypothetical protein | gp293.1 | - |
| gp059 | Hypothetical protein | gp248 | gp303 |
| gp094 | Hypothetical protein | gp219 | - |
| gp110 | Hypothetical protein | gp206 | - |
| gp123 | Hypothetical protein | gp194 | gp227 |
| gp130 | Hypothetical protein | gp187 | - |
| gp135 | Hypothetical protein | gp183 | - |
| gp150 | Hypothetical protein | gp168 | gp197 |
| gp177 | Hypothetical protein | gp145 | gp164 |
| gp199 | Hypothetical protein | gp124 | gp140 |
| gp270 | Hypothetical protein | gp068 | gp062 |
| gp279 | Hypothetical protein | gp059 | gp055 |
| gp325 | Hypothetical protein | - | gp115 |
| gp335 | Hypothetical protein | gp014 | gp122 |
| gp354 | Hypothetical protein | gp304 | gp376 |

To elucidate potential interaction partners of gp335, gp335-sfGFP was used as a bait, expressed in *P. aeruginosa*, and specifically pulled-down using anti-GFP antibody, followed by subsequent analysis via LC-MS/MS for protein identification. The sfGFP alone were used as an internal control for gp335-sfGFP and the proteins that were pulled down with the sfGFP were subtracted from the list to finalize the putative molecular interactors of the Churi gp335 protein **(Supplementary Table 4)**. Among 20 highly scored proteins, six of them were ribosomal proteins. These include 50S ribosomal proteins; L22, L28 and L4, and 30S ribosomal proteins; S9, S4, and S5. This result is consistent with our recent report demonstrating that gp014-phiKZ, the gp335-Churi homolog, interacts with the *P. aeruginosa* ribosomes^12^. This finding suggests the conserved function of these proteins in interacting with the host ribosomes. Additionally, we also found three putative interacting partners that possess high score above our selection criteria and are involved in bacterial RNA transcription **(Table 2 and Supplementary Table 4)**.

**Table 2.**
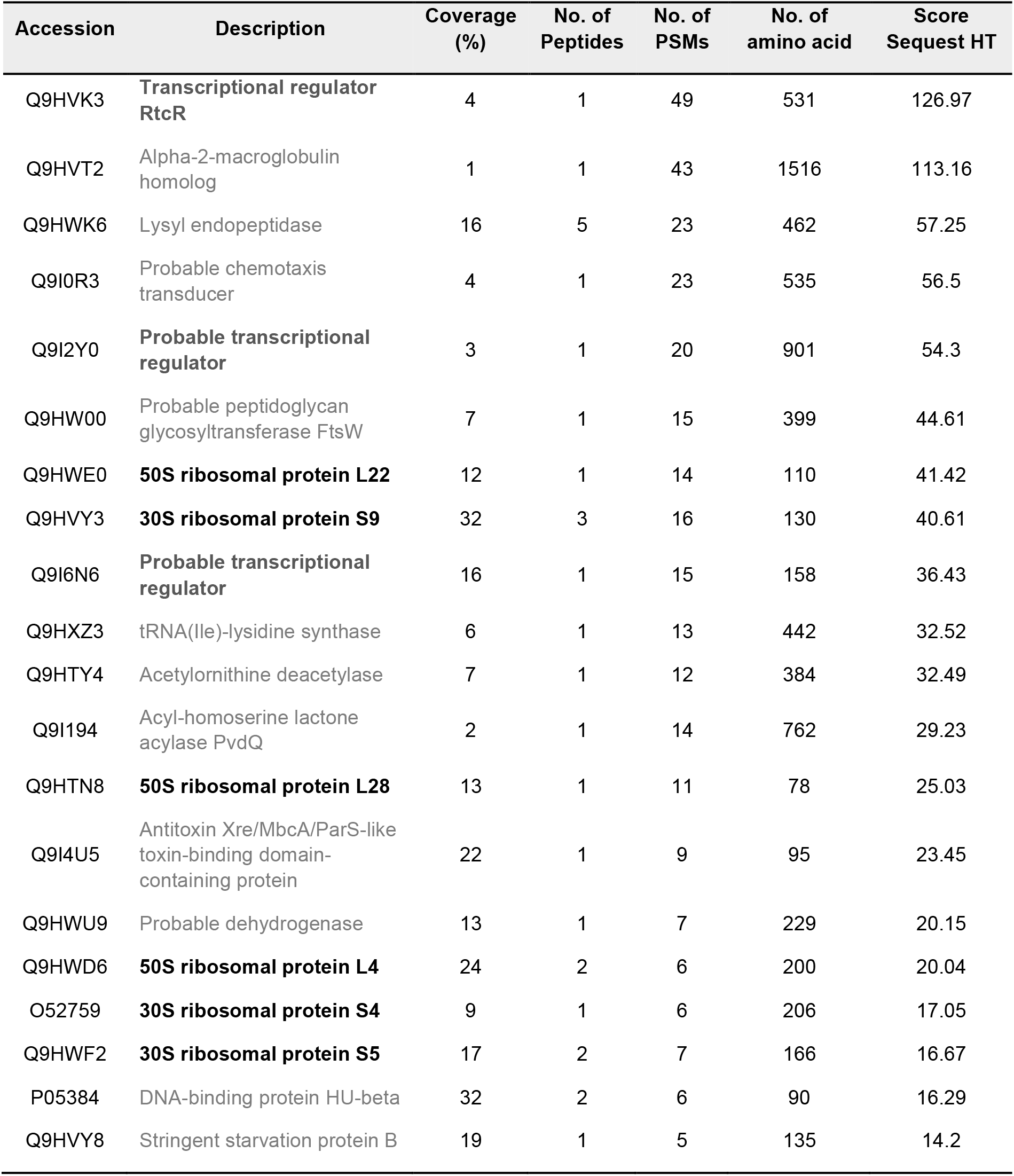
List of first twenty *P. aeruginosa* PAO1 proteins according to Sequest HT (abundant) score ranking found from co-immunoprecipitation (Co-IP) experiment with gp335-sfGFP.

### Gp335 does not trigger host cell morphological changes, but accumulates around the phage nucleus throughout Churi infection

Since some phage-derived proteins exhibiting antimicrobial properties have been shown to trigger morphological changes of the bacterial cells^16,17,29,30^, we then tested if the expression of gp335 at cytostatic level affects *P. aeruginosa* morphology at a single cell level. gp335-pHERD30T was transformed into the *P. aeruginosa* K2733 that is used for the microscopy experiment and the protein expression was induced with 0.8% arabinose. The result revealed that there was no such apparent change in cell morphology caused by the expression of gp335 alone **(Figure 3a)**. Both DNA and cell membrane are still intact and appear similar to the uninduced condition and the negative control (empty-pHERD30T) **(Figure 3a and Supplementary Figure 3)**.

**Figure 3.**
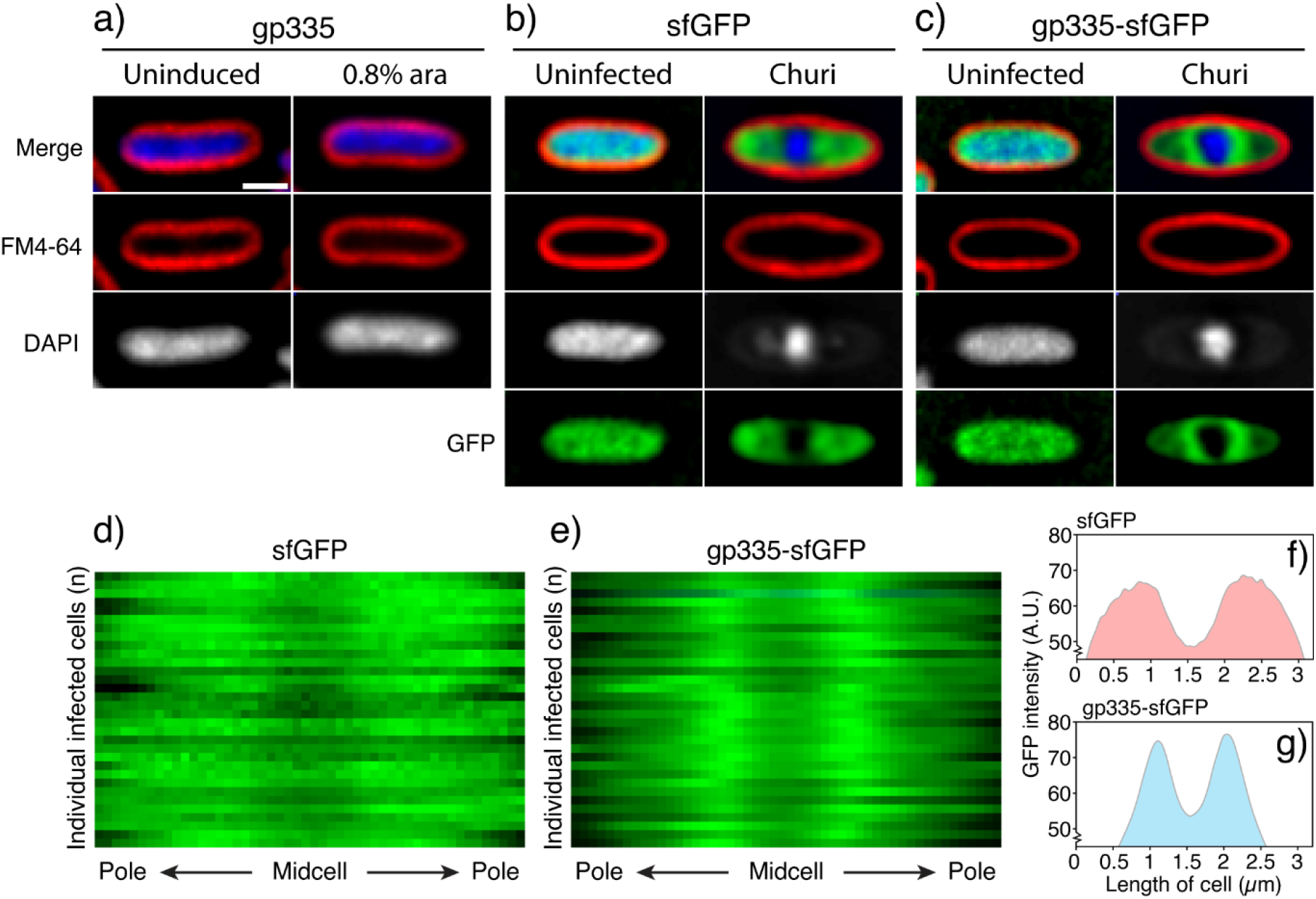
gp335 itself does not change host morphology while GFP-tagged gp335 accumulates close to phage nucleus during Churi infection. **a)** The fluorescence images of *P. aeruginosa* K2733 containing gp335-Churi when induced with or without arabinose. Localization of GFP-tagged proteins, induced with 0.2% arabinose when *P. aeruginosa* K2733 was infected with Churi at 90 mpi compared to uninfected control, **b)** sfGFP and **c)** gp335-sfGFP. Scale bar represents 1 µm. Kymograph shows the distribution of GFP intensity of individual infected cells (n = 32) at 90 mpi, **d)** sfGFP control and **e)** gp335-sfGFP. Distribution plots corresponding to the kymograph show the different pattern of GFP intensity between **f)** sfGFP control and **g)** gp335-sfGFP. A.U. means arbitrary units.

To further investigate if gp335 is spatially organized in the bacterial cells during the infection, we then created C-terminal fusion of gp335 with sfGFP and observed the protein localization in the cells. In the absence of phage, gp335-sfGFP is distributed uniformly in the cells similar to the control sfGFP alone **(Figure 3b, 3c, and Supplementary Figure 3)**, suggesting that gp335 is soluble in the cytoplasm. During infection at 90 mpi, gp335-sfGFP is excluded from the phage nucleus similar to sfGFP alone **(Figure 3b and 3c)**, suggesting that gp335 is not selectively transported into the phage nucleus. However, in addition to the localization in the cytoplasm, gp335-sfGFP accumulates around the phage nucleus appearing as a cloud-like pattern in the Z-section **(Figure 3c)**. Kymographs depicting 1-pixel horizontal slices from the central axis of 32 infected cells further illustrate that, unlike sfGFP control that is diffused from pole to pole and excluded from the phage nucleus **(Figure 3d)**, the GFP intensity of gp335-sfGFP is brightest at adjacent to the phage nucleus while is dimmer at the midcell where the nucleus localizes and the cell poles **(Figure 3e)**. Correspondingly, sfGFP intensity plot confirms the distribution of sfGFP along with cell length **(Figure 3f)** while the intensity plot of gp335-sfGFP exhibits narrow peaks beside the phage nucleus **(Figure 3g)**.

To determine the spatial-temporal dynamics of gp335 localization over time, we applied time series imaging on Churi-infected cells expressing gp335-sfGFP and visualized the protein localization as the infection progresses. We found that, at 15 mpi, the DAPI-stained early phage nucleus appears near the cell pole with gp335-sfGFP accumulated adjacent to it. As the nucleus moves towards the cell center at 30 mpi, the adjacent gradient of gp335-sfGFP moves along with it and remains associated with the phage nucleus. The cloud of gp335-sfGFP becomes denser and forms a ring-like morphology in the Z-section, encircling the phage nucleus at 60 mpi. This structure becomes larger in diameter as the phage nucleus grows in size from 60 to 120 mpi **(Figure 4a and Supplementary Figure 4)**. Later during infection, phage bouquets assemble as visualized by DAPI staining and these structures excludes gp335-sfGFP, similar to what is previously observed in other nucleus-forming phages **(Figure 4a)**^21^. Time-lapse imaging and fluorescence intensity plots of a single-infected cell over the course of the infection cycle confirms the spatial-temporal dynamic of gp335 that remains associated with the phage nucleus throughout the infection **(Figure 4b and 4c)**. Altogether with our finding that gp335 is an early-expressed protein and interacts with the host ribosomes, this finding further suggests that gp335 rapidly accumulates around the early phage nucleus and remains associated with it when it grows to serve a role in localized protein translation throughout the infection cycle.

**Figure 4.**
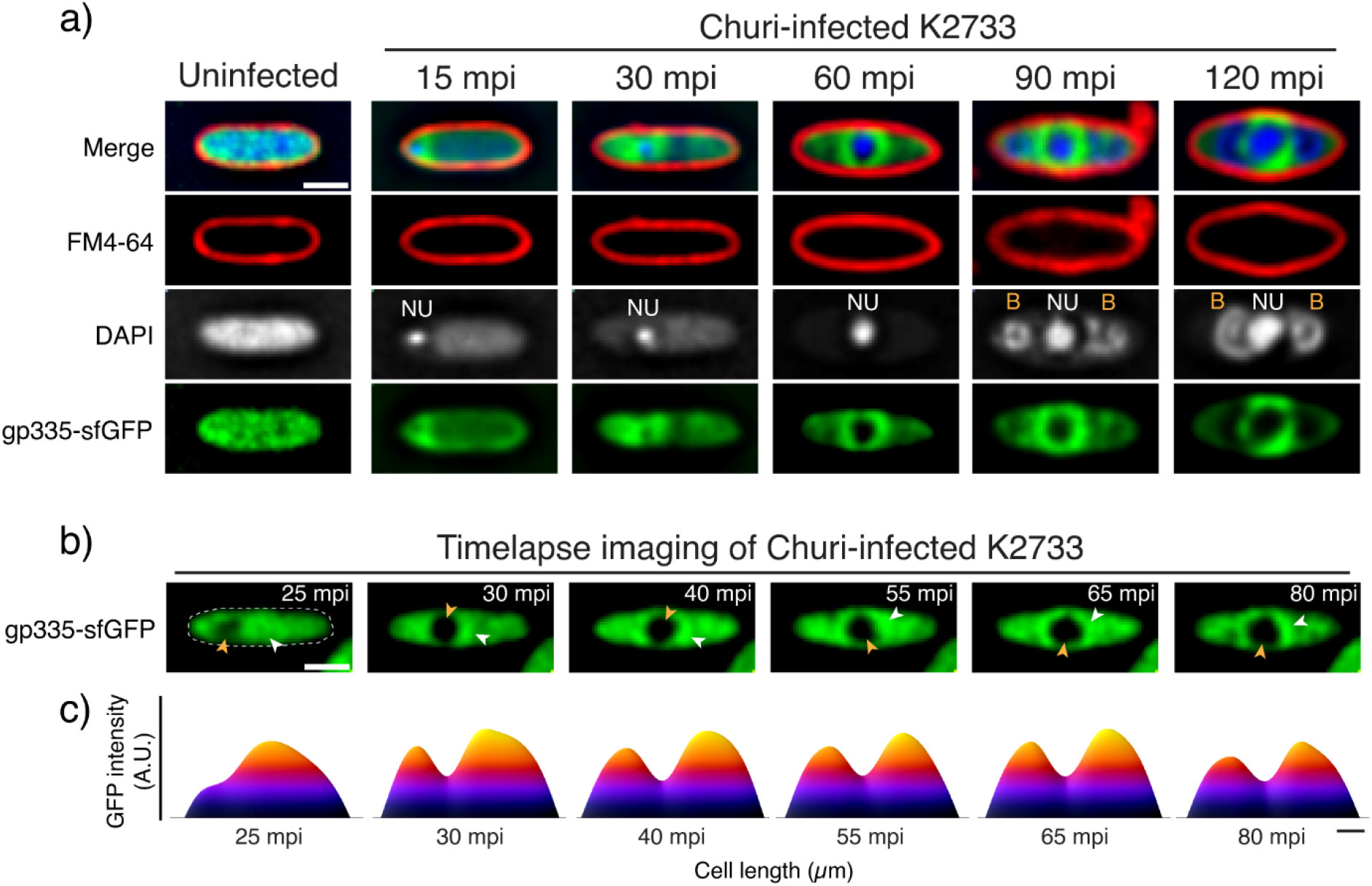
gp335 accumulation appears around phage nucleus at early infection and moves along with the nucleus as Churi infection progresses. **a)** Time-series images show the localization of GFP-tagged gp335 (green) during Churi infection against *P. aeruginosa* K2733 from early to late time point (minutes post-infection, mpi) compared to an uninfected cell. **b)** Time-lapse images show the single cell infection of Churi against *P. aeruginosa* K2733 expressing GFP-tagged gp335. Yellow arrowhead indicates the location of phage nucleus while the white arrowhead indicates the area of GFP with dense intensity. Scale bar represents 1 µm. **c)** GFP intensity heatmap of gp335 corresponding to the time-lapse imaging using raw images reveals the movement and accumulation of gp335 along with the Churi infection. Scale bar in **c)** represents 0.5 µm. A.U. means arbitrary units. NU and B are abbreviated from phage nucleus and bouquets respectively.

### gp335 is not essential for phage replication, but its mutant delays bacterial cell lysis

To determine if gp335 was essential for Churi replication, *Leptotrichia buccalis* CRISPR-Cas13a (Cas13) with a guide RNA targeting gp335 was used to select for naturally occurring mutant phage with gp335 mutated **(Figure 5a)**. Since Cas13 provides a strong selective pressure against the targeted sequence, only phage that have mutated that sequence can propagate. Otherwise, it would lead to abortive infection. If gp335 was essential, we would expect to only find mutants containing silent or missense mutations that might retain gp335 function. However, if gp335 was non-essential, nonsense mutations can be present. Our genomic sequencing of derived mutants revealed that gp335 was non-essential due to the presence of non-sense mutations early in the gene. A mutant named gp335G2-2 was selected for further study as it contained an inserted nucleotide, adenine (A) at position +41 base pair of gp335, which caused a frameshift mutation and an early stop codon. This shortened gene in the mutant is only 72 bp long compared to the wildtype gene (1,113 bp) and only the first 13 translated amino acids are similar to the wildtype since the following residues are translated out-of-frames until the amino acid at position 23 **(Figure 5b)**. In contrast, we have previously used this method to select for mutants of the essential gene gp069-phiKZ, in which only missense or silent mutations were found^31^. Therefore, the nonsense mutation obtained in gp335G2-2 that subsequently leads to a frameshift and premature stop codon causing large change in gp335-Churi product indicates that the gene is not essential for Churi viability. We then investigated whether the mutation in gp335 is involved in the progeny production by evaluating the phage titer of the gp335G2-2 mutant compared to that of the wildtype. The result revealed that although the phage titer of the gp335G2-2 mutant is lower than that of the wildtype (gp335WT), they are not statistically different **(Figure 5c)**. To further examine if the loss-of-function of gp335 impairs the phage fitness that might lead to a prolonged latent period, we performed the single-step time to lysis to identify a difference in latent periods between the wildtype and gp335G2-2 mutant. This technique is highly accurate in measuring the bacterial cell lysis as it uses a cell-impermeable fluorescent DNA staining dye which, upon cell lysis, it can stain the released nucleic acid and rapidly increases in brightness for detection^32^. We found that, as determined by the point at which 50% of maximal fluorescence is reached, wildtype Churi lyses the cell on average at 78 mpi while the gp335G2-2 mutant lyses at 82 mpi **(Figure 5d)**, indicating the delay of bacterial cell lysis when infected with the mutant. Complementing the infection by Churi gp335G2-2 with the wildtype gp335 protein expressed from pHERD30T after induction can rescue the replication cycle of the mutant by decreasing the latent period of the mutant up to 4 minutes compared to the uninduced control, further suggesting that even though gp335 is not essential for phage replication, it serves a role in maintaining the phage fitness to shorten the latent period of the replication cycle **(Figure 5e)**. Uninduced or induced empty-pHERD30T with arabinose had no effect on the time to lysis of the gp335G2-2 mutant **(Figure 5f)**. To further investigate whether the inactive mutant gp335 expressed by Churi gp335G2-2 can retain the localization, we created the C-terminal fusion of the mutant gp335 with sfGFP and expressed it to visualize the localization of the mutant during infection. The result revealed that, unlike the wildtype protein, no ring-like pattern was observed around the phage nucleus in the mutant **(Figure 5g)**, suggesting that the mutant gp335 loses an ability to accumulate around the phage nucleus for localized protein translation, thereby contributing to the delay of bacterial cell lysis.

**Figure 5.**
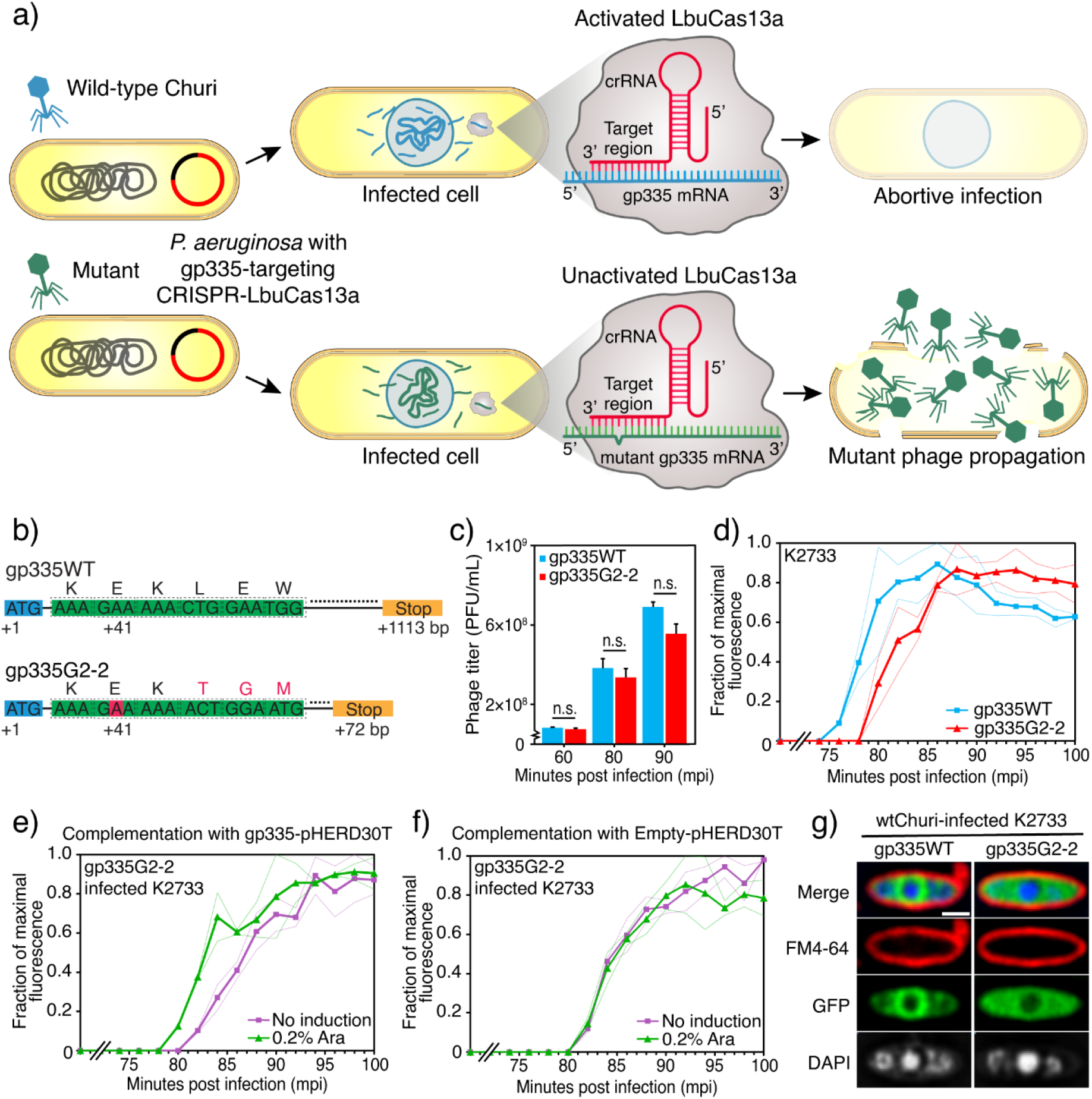
gp335-mutant Churi slightly delays in cell lysis and complementation of gp335 improves time to lysis of the mutant phage. **a)** The schematic mechanism of mutant phage isolation with LbuCas13a. An anti-sense guide RNA complementarily pairs with the mRNA transcript of gp335. This activates the Cas13a that leads to abortive infection of Churi bearing the wildtype gp335. The mutant phages, that already exist as the sub-population escape this selection and become major or dominant populations. **b)** Variant calls on gp335 from whole genome sequence of wild type (gp335WT) and mutant-gp335G2-2 Churi shown as a gene diagram indicate a nucleotide insertion at early sequence of gp335 (red box). This leads to frameshift mutation (red alphabets) and stop codon at +70 bp. Numbers represent nucleotide position in the gene. Orange box is a stop codon. **c)** Titer count (PFU/mL) in different minutes post infection between wild type (gp335WT) and gp335G2-2 mutant Churi as determined by spot test on *P. aeruginosa* K2733 lawns. N = 3 per condition. N.s. means non significance at *p* ≤ 0.05. **d)-f)** Single-step time to lysis represents real-time bacterial cell lysis caused by the phage infection. Phage progenies, that release from infected cells, are detected by SYTOX green. **d)** Lysis pattern of wildtype Churi compared to mutant gp335G2-2. Complementation experiment shows lysis time of **e)** mutant gp335G2-2 Churi against *P. aeruginosa* expressing gp335 when gp335 was induced with or without 0.2% arabinose while **f)** empty-pHERD30T was used as a control to ensure that arabinose does not affect the lysis of Churi. Solid lines depict the mean across n = 6 replicates. Faint lines show mean +/- standard deviation. **g)** Fluorescence images between *P. aeruginosa* K2733 cells that expressed either wild-type gp335-sfGFP or gp335G2-2-sfGFP when they were infected with wild-type Churi at 90 mpi. Scale bar represents 1 µm.

## Discussion

Jumbophages have been of interest recently, not only because of their intricate replication strategies^31,33–36^, but also for their therapeutic potential and biocontrol application^37–41^. This is partially due to the presence of a significant number of uncharacterized genes that facilitate the invasion and regulation of them in their bacterial host^22,23,42^. We initiated this study focusing on isolating a jumbophage and successfully obtained a new *P. aeruginosa* jumbophage, Churi. The genomic analysis of Churi revealed that it is closely related to phiKZ, representing a variant of the over 30 different phiKZ-like phages that have been reported in the Genbank database to date. Churi contains homologs of ChmA (a major component of the phage nuclear shell), PicA (involved in protein import into the phage nucleus), and PhuZ (orchestrating the nuclear position and capsid trafficking). These homologs are conserved among nucleus-forming phages^23,31^. Additionally, the morphology and protein localization of Churi-infected cells are consistent with those of other nucleus-forming phages, as evidenced by the microscopy data **(Figure 1d and Supplementary Figure 1)**. Altogether, genomic analysis and microscopy indicate that Churi is a previously unidentified member of the phiKZ-like phages within the *Chimalliviridae* family^23^. Despite the fact that the genomic contents of Churi are largely similar to those of phiKZ, we have observed that not all genes of Churi are conserved in phiKZ. Approximately 4% of all genes are low-conserved, with a percent identity between Churi and phiKZ that is less than 80%. These proteins include gp056-phiKZ and gp283-Churi (HNH nuclease; 35.1% identity), gp094-phiKZ and gp238-Churi (the internal head protein; 40% identity), gp303-phiKZ and gp355-Churi (hypothetical protein; 70.1% identity), and gp299-phiKZ, which is previously found to interacting with 50S ribosomal subunit^12^, and gp358-Churi (hypothetical protein; 69.5% identity). Some genes are uniquely found in the phage. For example, gp001, gp208, gp209, gp325, and gp340 (Methyltransferase type 11) are uniquely present in Churi but absent in phiKZ, while gp072 (HNH endonuclease) and gp296 of phiKZ are absent from the Churi genome. These distinct genotypes suggest that there may be some interspecies differences contributing to the unique phage-host interplay between these bacteriophages. In addition to the differences at their genomic level, the replication machinery within the host is also somewhat distinct. Even though Churi is highly closely related to phiKZ, the frequency of bouquet formation during late infection of Churi is more similar to phiPA3, which is more distantly related than to phiKZ. Bouquets are observed in Churi around 70% of all infected cells at 90 mpi **(Supplementary Figure 1**), compared to 20% in phiKZ and 80% in phiPA3^21^. Further investigation into the difference between genomes’ Churi and phiKZ and their encoded products will shed light on the factors necessary for jumbophage bouquet assembly.

Recently, a number of studies have reported that bacteriophages express a variety of proteins during early infection, which disturb cellular processes and restrict bacterial growth^14–20^. For instance, early-expressed genes from phiKZ (gp058, gp124, and gp287) were also identified to bind to bacterial DnaX and FtsZ; however, these proteins do not inhibit the growth of *P. aeruginosa*^15^. Thus, we sought to identify early jumbophage proteins that inhibit the growth of the host bacterium. Our proteomic studies with Churi revealed early expressed gene candidates, among which we identified gp335 as a growth inhibitor of *P. aeruginosa*. The closest homolog of gp335-Churi, gp014-phiKZ, has also recently been discovered to be expressed early during phiKZ infection^12^. This finding is consistent with our gp335 result in Churi. Specifically, our results further demonstrated that, similar to gp335-Churi, gp014-phiKZ possesses a potent inhibitory effect on the growth of *P. aeruginosa* **(Figure 2)**. It is not surprising that the activity of both gp335-Churi and gp014-phiKZ is similar, since the amino acid sequence of gp335-Churi is highly identical (99.73%) to that of gp014-phiKZ **(Supplementary Figure 5 and Supplementary Table 5)**. On the other hand, gp122-phiPA3, whose amino acid sequence is only 28.41% identical to that of gp335-Churi, does not appear to have this antimicrobial property **(Figure 2 and Supplementary Table 5)**. We previously elucidated that gp014-phiKZ requires certain conserved amino acids for its complete interaction with ribosomes, including R10, K15, W18, R23, K152, R262, and K263^12^. Our multiple alignment of gp335 and its homologs, gp014-phiKZ and gp122-phiPA3, revealed that even though most amino acid residues are conserved among all 3 phages, suggesting their role in binding with ribosomes, K15, W18, and K152 (phiKZ) are only conserved in gp335-Churi but not in gp122-phiPA3 **(Supplementary Figure 5)**. It is important to note that homologs in nucleus-forming phages can still perform comparable functions, despite sharing a very low sequence identity. For example, the PhuZ proteins in nucleus-forming phages (201phi2-1, phiKZ, and phiPA3) behave the same dynamic function during phage infection despite the fact that their amino acid sequence similarities are only 30-40%^43,44^. Based on this evidence, we remain convinced that gp122-phiPA3 may possess a central function that is comparable to that of Churi and PhiKZ, albeit with a less potent inhibitory effect on bacterial growth. Further investigation into gp122-phiPA3 on these less conserved residues (K15, W18, and K152) might shed light on the growth inhibitory activity of this protein. It is also worth noting that, gp110-Churi **(Table 1)**, a homolog of gp206-phiKZ that has been reported to co-sediment with the 50s ribosomal subunit^12^, unlike gp335-Churi, does not exhibits an inhibitory effect against *P. aeruginosa* **(Figure 2a)**. This finding emphasizes the unique antimicrobial properties of gp335-Churi.

Throughout the Churi infection, gp335 is spatiotemporally localized adjacent to the phage nucleus. At 15 mpi, the gradient of gp335 is formed and accumulated at the immature phage nucleus, which is near the cell pole. Our finding corresponds to previous findings that has demonstrated that gp014-phiKZ is rapidly recruited to an early phage infection (EPI) vesicle that assembles at the early stage of the nucleus-forming phage infection cycle (10 mpi), likely playing a key role in the translation of early-expressed phage proteins^45,46^. Throughout the infection, gp335 always accumulates around, but not inside the phage nucleus. This indicates that gp335 is not involved in DNA replication or transcription^47^ but is likely associated with certain activities that must be proximal to the phage DNA on the nuclear shell, such as the translation of mRNA exported from the phage nucleus. Our Co-IP results demonstrate that gp335-Churi interacts with host ribosomes. This data agrees well with our previous study of gp014-phiKZ, which has been identified as a ribosome-binding protein. gp014-phiKZ was found to co-sediment with the host large-subunit ribosome in the Grad-seq experiment, and our cryoEM result further verified the binding of gp014-phiKZ to the bacterial ribosome^12^. However, the ribosomal proteins that are identified in this study are not comparable to the ribosomal proteins that were studied in phiKZ^12^. Additionally, we also identified host transcriptional regulators as putative molecular interactors of the Churi gp335, implying that gp335 might also interfere with the transcription of *P. aeruginosa* gene during Churi infection. Further investigation into the function of gp335 in transcriptional interference will be required.

Cas13 selection of gp335-null Churi mutants revealed that gp335 is non-essential for phage propagation. However, the loss-of-function of gp335 can cause a delay in the Churi lytic life cycle but can be rescued by exogenous expression of the wildtype gp335. This finding suggests that gp335 is important for efficient Churi propagation within the host possibly via maintaining phage fitness. It is worth mentioning that, the negative impact of losing gp014-phiKZ was not quite apparent in the lab strain, *P. aeruginosa* PAO1, but its mutant has a more obvious negative impact on phage propagation when infecting specific clinical isolates of *P. aeruginosa*^12^. To gain insights into the phage interplay with diverse bacterial hosts, further investigation into the effect of gp335-Churi in other *P. aeruginosa* strains might be necessary. Overall, gp335-Churi is clearly beneficial for the efficient completion of the phage lytic cycle. This has evolutionary implications since faster completion of the lytic cycle can provide phages with a selective advantage over other slower competitors^48,49^.

## Materials and methods

### Phage isolation, purification, and propagation

A soil sample collected from Chulalongkorn university, Bangkok, Thailand was enriched by adding LB medium, 0.5 mM CaCl_2_, and overnight culture of *P. aeruginosa* PAO1. The mixture was incubated at 37°C for 48 hours with shaking at 200 rpm. After the incubation, the phage-enriched culture was centrifuged at 8,500 rpm, 4°C for 10 minutes and filtered through 0.45 μm filter. Double-layer agar (DLA) plates were prepared by mixing the 10 µL of enriched filtrates, 100 µL of *P. aeruginosa*, and 5 mL of 0.35% LB agar (Top agar). Then, the mixtures were transferred onto LB plates (Bottom agar). After incubation, single plaques were selected and kept in SM buffer. Plaques with same morphology were selected at least 3-5 repeats to ensure that the isolated phages were purified. To propagate high titer bacteriophage, phage-containing SM buffer was diluted into the suitable dilution. At the concentration that web lysis in which plaques merged to another throughout the plate, appeared, the plates were soaked with 5 mL of SM buffer for 5 hours. After that, the phage lysate was collected and centrifuged at 8,500 rpm, 4°C for 10 minutes before filtration with 0.45 µm. The phage lysate was kept at 4°C, and phage titer was counted by spot test^28,43^.

This work has been reviewed and approved by Chulalongkorn University-Institutional Biosafety Committee (CU-IBC) in accordance with the levels of risk in pathogens and animal toxins listed in the Risk Group of Pathogen and Animal Toxin (2017) published by Department of Medical Sciences (Ministry of Public Health), the Pathogen and Animal Toxin Act (2015) and Biosafety Guidelines for Modern Biotechnology BIOTEC (2016) with approval number: SC CU-IBC-028/2020 Ex1.

### One-step growth curve

*P. aeruginosa* PAO1 in exponential phase (OD600∼0.4) was infected with phage Churi at MOI of 0.01 for 15 min at 37°C. Then, the cells were centrifuged at 8,500×g for 2 minutes. The pellet was washed with LB broth before centrifugation at 8,500×g for 2 minutes. After discarding supernatant, the pellet was resuspended with 10 mL of LB broth and incubated in a shaker at 250 rpm, 37°C for 2 hours. Every 10 minutes including time point 0, the phage-bacterial mixture was collected and filtered with 0.45 µm filters. The filtrates were counted by spot titer assay. The PFU/mL will be calculated and plotted against minutes post infection (mpi). The latent period was observed. Likewise, phage burst size was calculated from the plotted curve as particles/cell^37,50^.

### Transmission electron microscopy (TEM)

Phage Churi lysate was precipitated with 10% (w/v) of polyethylene glycol and 1 M NaCl overnight. The precipitated phage was centrifuged at 8,500 rpm for 10 minutes and resuspended with SM buffer. The phage was negatively stained with 2% uranyl acetate on a carbon-coated grid and the morphology of phages were visualized by transmission electron microscope (HITACHI model HT7700). Then, the phage classification was based on their structure according to ICTV classification observed under TEM.

### Phage genomic DNA extraction

Genomic DNA of phage Churi was extracted using phenol-chloroform–isoamyl alcohol method. Briefly, the phage lysate with at least 10^9^ PFU/mL was mixed with 10% (w/v) polyethylene glycol and 1M NaCl overnight to precipitate the phage particles. Phage precipitate was centrifuged at 8,500 rpm for 10 minutes^51^ before resuspended with SM buffer. After that, host DNA and RNA were digested with 10U of DNase I and 0.1 mg/mL of RNase A at 37°C. The DNase and RNase were inhibited by 20 mM EDTA. Then, phage capsid was digested by 0.5 mg/mL of proteinase K and 0.5% sodium dodecyl sulfate (SDS). After the incubation at 55°C for 1 hour, Phenol-Chloroform–Isoamyl alcohol (25:24:1) (Sigma Aldrich, Switzerland) was added to the mixture at equal volume. The mixture was gently mixed before centrifuged at 15,000×g for 10 minutes and the top aqueous layer was collected. The steps of Phenol-Chloroform–Isoamyl alcohol and centrifugation were repeated before treated with 1/10 volume of 3 M sodium acetate and 2 volumes of cold absolute ethanol. The genomic DNA was allowed to precipitate at -20°C for 3 hours followed by centrifugation at 21,000×g for 20 minutes. The DNA pellet was washed with 70% ethanol, centrifuged at 15,000×g for 5 min, and the pellet was air-dried until the remaining ethanol evaporated. The phage DNA was dissolved with TE buffer. DNA quality and quantity were analysed using a NanoDrop 2000 spectrophotometer (Thermo Scientific)^28,52^.

### Whole genome sequencing (WGS), genome analysis and comparison

Whole genome sequencing was performed using the MiSeq (Illumina). FASTQ files with 2,110,430 reads of Churi genome derived from the sequencing were assembled with Unicycler version 0.4.8^53^. The single contig with coverage over 1,500x was then annotated with DNA master version 5.23.6 (Glimmer 3.02 and GeneMark HMM). Functional prediction was manually performed using viral genomes available in the NCBI database, NCBI conserved domain, Phyre2 were additionally used for the annotation. The phage genome map was constructed and visualized using Artemis: DNAPlotter version 18.1.0. Genetic relation and genome organization between phage Churi and other nucleus-forming jumbophages as followings phiKZ (NC_004629.1), phiPA3 (NC_028999.1) and 201phi2-1 (NC_010821.1) were compared by Easyfig software version 2.1^54^. Pairwise intergenomic similarities between related phage genomes were performed using the Virus Intergenomic Distance Calculator (VIRIDIC; http://rhea.icbm.uni-oldenburg.de/VIRIDIC/)^24^. The VIRIDIC algorithm relies on ICTV criteria to calculate intergenomic similarities among bacteriophages for classification.

### Proteomics study of phage Churi during infection

We performed proteomics study in different infection conditions. First is the infection on solid medium. Briefly, mid-log phase or OD_600_∼0.6 (∼6×10^8^ CFU/mL) of *P. aeruginosa* PAO1 was centrifuged at 4,500 rpm for 10 minutes. The pellet was resuspended with LB broth before spreading on LB plates and incubated for 5 hours. Then, high titer phage Churi was gently spread onto the plate with MOI of 1. At 15 minutes post infection, the bacteria-phage mixtures were flooded with LB broth and gently scraped with L-shape spreader before transferred into microcentrifuge tubes. The following steps were performed on ice or 4°C. After centrifugation at 15,000xg for 1 min, the supernatants were discarded and resuspended with 1X lysis buffer (2% SDS, 3 mM IAA, 1X RIPA buffer, and 1X Protease inhibitor cocktail). Samples were sonicated at 21% amplitude for 10s with 1s pulse on/off and pelleted at 15,000×g for 1 min to remove the cell debris and intact cells^47^. The supernatants were immediately kept at -80°C before. For the infection in liquid culture, log phase *P. aeruginosa* PAO1 was mixed with phage Churi at MOI of 15^55,56^. At 15 minutes post infection, the mixtures were rapidly cooled on ice and collected at 5,000 rpm, 4°C for 5 minutes. The pellets were washed three times with ice-cold 1X PBS and resuspended in 500 µL of 1X lysis buffer. The samples on ice were sonicated at 21% amplitude for 3 minutes with 5s pulse on/off before centrifugation at 15,000×g for 1 minute to remove the cell debris and intact cells. The samples were kept at -80°C. Both solid and liquid infections were performed in duplicate. To analyze the phage Churi proteins expressed at 15 minutes post infection, the protein concentration was measured before injected into LC-MS/MS. The LC-MS/MS analysis was performed at Department of Biochemistry, Faculty of Science, Mahidol university, Bangkok. Phage structural proteins were subtracted from all peptide signals that are above the cutoff value. Churi proteins found to be expressed in both solid and liquid-mediated infection were selected for further experiments.

### Screening of phage-encoded growth inhibitory proteins

The selected non-structural genes of Churi that expressed during the early infection were cloned into *E. coli*-*Pseudomonas* shuttle vector pHERD30T^47^ using Gibson assembly. The correctness of recombinant plasmid was verified by Sanger sequencing. Then, the construct was electroporated into competent cells of *P. aeruginosa* PAO1. The recombinant cells were cultured in LB with high gentamicin selection (300 µg/mL) to keep the plasmid. To test the growth inhibition activity of Churi proteins, 200 µL of LB broth containing gentamicin with different concentrations of arabinose or without the arabinose was dispensed into 96 well-plates. Overnight cultures of recombinant *P. aeruginosa* were diluted until OD_600_ ∼0.2 and further 10-fold diluted. One microliter of each recombinant culture was added into the wells in triplicate. The 96-well plates were wrapped with parafilm to prevent evaporation and incubated at 37°C for 18 hours. After incubation, the culture from each well was 10-fold serially diluted and 5 µL of each diluted culture was spotted onto LB agar containing gentamicin. The plates were incubated at 37°C for 18 hours and colony-forming units were then calculated. Empty-pHERD30T and gp10 (JJ01)-pHERD30T were used as a negative control and positive control respectively.

Real-time activity against *P. aeruginosa* of phage proteins was also observed by measuring optical density (OD_600_) of *P. aeruginosa* that expressed phage proteins. Bacterial cells were cultured overnight before the OD_600_ was adjusted into ∼0.2, then 1 µL of each cultured were transferred into 96-well plates containing LB with gentamicin and different concentrations of arabinose from 0.2, 0.4, and 0.8%. The OD_600_ was measured every 20 minutes after gene induction for 10 hours at 37°C.

### Pulldown experiment for protein-protein interaction analysis

Overnight culture of gp335-sfGFP *P. aeruginosa* PAO1 was diluted into OD_600_∼0.1. Then, the cells were further grown in 150 mL of LB broth supplemented with 25 µg/mL of gentamicin and 0.4% arabinose until OD_600_∼0.3 followed by centrifugation at 4,000 rpm, 4°C, for 10 minutes. After that, the pellets were transferred into microcentrifuge tubes and then resuspended with 1 mL of lysis buffer (0.1% lysozyme, 25 mM Tris; pH 7.5, 150 mM NaCl, 4 mg/mL lysozyme, 20 µg/mL DNase I, 2X cOmplete^Tm^ protease inhibitor cocktail, and 0.4 mM PMSF)^57^. The reaction tubes were incubated at 37°C for an hour. The cells were further disrupted using probe tip sonication at a frequency of 20 kHz and 80% amplitude for 2 seconds on, and 8 seconds off at the total of 20 seconds before centrifugation at 14,000xg, 16°C for 20 minutes. The supernatants were collected, and protein concentration was measured by BCA protein assay. The protein concentration was then adjusted to 1 µg/µL with the 1xPBS+0.1%Tween20. Ten microliters of slurry protein A beads were added to the protein solution for 1 hour at 37°C to remove non-specific binding. The mixtures were transferred to centrifugal filter (Thermo Fisher) and the flow-through (FT) was collected. Anti-GFP antibody (Chromotek) (25 µL) was mixed with the FT and incubated at 8°C for 16 hours before adding 25 µL of protein A beads. The mixtures were incubated at 25°C for 3 hours and then transferred to centrifugal filter (Thermo Fisher). The flow-through (FT) was discarded at this time. The beads were washed with 400 µL of 1xPBS+0.1% Tween20 for three times, and then washed with 400 µL of 1xPBS for four times. The beads were collected and subjected to on-bead digestion. Briefly, beads were resuspended in 0.15% RapiGest SF Surfactant (Milford, MA, USA), 10 mM NaCl, and 10 mM ammonium bicarbonate. The mixture was subjected to probe tip sonication at a frequency of 20 kHz and 80% amplitude for 2 seconds. Subsequently, 2 mM of TCEP was added to the samples and incubated at 90°C for 15 minutes. After that, the samples were cooled down, and 10 mM IAA solution was added. The samples were incubated in the dark at room temperature for 45 minutes. After that, 50 ng/µl of trypsin in ammonium bicarbonate at a 1:50 w/w ratio was added, and the samples were incubated at 37°C for 4 hours. Lastly, 1% formic acid at a 1:10 v/v ratio was added to terminate the reaction, and the tryptic peptides were lyophilized before LC-MS/MS analysis.

### LC-MS/MS setting and protein searching for protein identification

The LC-MS/MS spectrum data were collected in the positive mode with an Orbitrap HF mass spectrometer combined with nano-LC system equipped with an EasySpray C18 column (Thermo Scientificɛ ES903; 75 μm x 50 cm, 2.0 µm) using previous protocol with minor modifications^58^. Briefly, mobile phase A consisted of 0.1% formic acid in water and mobile phase B consisted of 100% acetonitrile with 0.1% formic acid. Separation was conducted with a linear gradient of 5%–45% mobile phase B at a constant flow rate of 300 nL/min over a period of 115 min. The tryptic peptides were analyzed by applying a data-dependent acquisition method, followed by a higher-energy collisional dissociation. Full scan mass spectra were acquired at an *m/z* ratio of 400 to 1600 with an AGC target set at 3×10^6^ ions, a resolution of 120k, and injection time was 60 ms. MS/MS scanning was initiated when the automatic gain control target reached 3e^6^ ions and, a resolution of 60k, and injection time was 100 ms. Isolation windows were 1.6 *m/z*. Xcalibur software (Thermo Fisher) was utilized to automatically collect the mass spectrum. Raw LC–MS/MS files underwent analysis using the Proteome Discoverer with SEQUESTɛ HT algorithm (Thermo Fisher), referencing the in-house protein database, adhering to specific criteria: strict trypsin specificity, up to two missed cleavages, a fixed carbamidomethyl modification on cysteine (+57.0215 Da) and variable modification on methionine (+15.9949). The relative protein abundance was standardized using software’s normalization algorithm. Proteins that had a FDR value< 1% were selected for further analysis.

### Single cell infection and fluorescence microscopy

To observe Churi infection morphology under fluorescence microscopy, ¼LB containing 1.2% agarose pads on concavity slides were prepared. Each pad contains 1 µg/mL FM4-64 for cell membrane staining, and 1 µg/mL DAPI for DNA staining^43,59^. Colonies of *P. aeruginosa* K2733 were resuspended in ¼ LB broth before 5 µL of the suspension was inoculated onto the agarose pad. K2733 strain is derived from PAO1 that multiple efflux pumps are knockout (ΔmexB, ΔmexX, ΔmexCD-oprJ, ΔmexEF-oprN) and is used in the microscopy experiment^44^. This improves cell staining with fluorescent dyes while identical results were obtained in both PAO1 and K2733^44^. The slides were then incubated in a humid chamber at 30°C for 3 hours. After that, 5 µL of high titer Churi lysate was added onto the pads. A cover slip was placed on the pad before fluorescence microscopy. For localization profiling, recombinant *Pseudomonas* cells that contain fluorescence tagged version of Churi proteins were grown on the agarose pads, supplemented with the suitable concentration of arabinose to induce phage gene expression for 3 hours at 30°C in the humid chamber prior to Churi infection. To observe the effect of gp335-Churi on bacterial cell morphology, overnight cultures of *P. aeruginosa* K2733 carrying gp335-, sfGFP-, and empty-pHERD30T were inoculated into LB broth, supplemented with gentamicin and different concentrations of arabinose. The cultures were rotated on a roller at 37°C for 3 hours. After incubation, the cells were centrifuged at 8,000 rpm for 1 minute. The pellets were resuspended and mixed with FM4-64 and DAPI mixture before added onto the agarose pads. The DeltaVision Spectris Deconvolution Microscope (Applied Precision, Issaquah, WA, USA) was used to visualize the cells. For static images, the cells were imaged for at least five stacks from the middle focal plane with 0.15 mm increments in the z axis and, for time-lapse imaging, the cells were imaged from a single stack at the focal plane with the ultimate focusing mode. Microscopic images were further processed by the deconvolution algorithm in the DeltaVision SoftWoRx Image Analysis Program^47^.

### Generation of gp335 mutant Churi with Cas13a

Plasmids were synthesized and cloned by Genscript (USA). The vector for all *P. aeruginosa* plasmids was pHERD30T (**Supplementary Table 6**). The plasmids were transformed at 2.5 kV electroporation into K2733 competent cells. *P. aeruginosa* K2733 that contains Cas13a expression vector pHERD30T-LbuCas13a with a guide targeting gp335-Churi were grown to OD_600_∼0.5-0.8 in LB containing 15 μg/mL gentamicin and 0.2% arabinose. One-hundred microliter of the culture was infected with high titer Churi lysate and mixed with 5 mL of LB 0.35% agar containing 0.2% arabinose at 55°C. Lawns of *P. aeruginosa* expressing gp335-targeting Cas13a were grown on LB plates containing 15 μg/mL gentamicin. Single plaques of Churi mutants escaping from Cas13a activity were selected and streaked to purify at least three times using bacteria expressing gp335-targeting Cas13a as the host^31^. Finally, the targeted region on Churi genome was amplified by PCR and the amplicon sequenced by Sanger sequencing to identify mutations and confirmed by whole genome sequencing. The phage lysate samples were sequenced by the SeqCenter (Pittsburgh, USA). After genomic extraction, whole genome sequencing was performed on an Illumina NovaSeq 6000 sequencer in one or more multiplexed shared-flow-cell runs, producing 2x151bp paired-end reads with coverage at 2,618. Demultiplexing, quality control and adapter trimming was performed with bcl-convert1 (v4.1.5). The variant calling was performed using reference genome sequence information of Churi (Accession No. OM718002.1).

### Single Step Time-to-Lysis

Single step time to lysis was applied to measure the effect of gene mutation on mutant Churi lysis time point in real time^32^. Briefly, *P. aeruginosa* strain PAO1-K2733 was grown in LB broth at 37°C to early-log phase (OD_600_ ∼0.25-0.3). High titer wild-type or mutant Churi was mixed with the culture at MOI of 5 and incubated at 30°C for 20 minutes before adding 5 µM SYTOX Green Nucleic Acid Stain (Thermo Fisher). Next, 200 µL of each phage-bacteria mixture was transferred into a black-walled, clear-bottom 96-well plate (Costar). Fluorescence and optical density measurements were performed in a microplate reader (Tecan Infinite M Plex) at 30°C. Fluorescence measurements were taken with the following settings: λ-excitation = 504 nm; λ-emission = 537 nm; gain = 25; flashes per well = 5. Optical density measurements were taken with the following settings: λ-excitation = 600 nm; flashes per well = 5. Measurements were taken every 2 minutes for 45 cycles, to a maximum of 120 minutes post-infection. The plate was shaken for 20 seconds and waited for 20 seconds before measurements. Data analysis was done with Microsoft Excel. For each replicate, background fluorescence was subtracted, and values were divided by the maximum fluorescence signal as a fraction of maximal fluorescence equal to 1. Negative values due to photobleaching of the background prior to lysis were set to zero. The mean +/- the standard deviation (SD) across replicates was plotted and average time-to-lysis determined as the time point at which the mean fraction of maximal fluorescence reached 0.5^31^. For complementation experiment, *P. aeruginosa* K2733 strain containing gp335-pHERD30T was incubated with 0.2% or without arabinose at 37°C before infected with gp335G2-2 mutant Churi.

## Data availability

The nucleotide sequence of the *Pseudomonas* phage Churi genome was deposited in the GenBank database with the accession number: OM718002.1. All data supporting this article are available in the article and/or the Supplementary data. If there are any special requests or questions for the data, please contact the corresponding author (VC).

## Acknowledgements

We acknowledge the support for scientific instruments by the Japan Science and Technology Agency (JST)/Japan International Cooperation Agency (JICA), Science and Technology Research Partnership for Sustainable Development, SATREPS JPMJSA1806 (VC and PN). We also thank Mahidol University Frontier Research Facility (MU-FRF) for Instrumentation Support for DeltaVisionTM Ultra at Mahidol University for the fluorescence microscopy imaging.

## Author contributions

WW, CA, PN, JP, and VC: conceptualization. WW, SK, PN, JP, and VC: methodology. WW, SK, CA, PN, JP, and VC: investigation. WW, SK, PN, JP, and VC: formal analysis. WW, SK, PN, JP, and VC: visualization. WW and SK: validation. WW, SK, PN, JP, and VC: writing of the original draft. All authors: reviewing and editing. PN, JP, and VC: project administration. CA, PN, JP, and VC: resource. PN, JP, and VC: supervision. CA, PN, JP, and VC: funding acquisition.

---

**Supplementary Figure 1.**
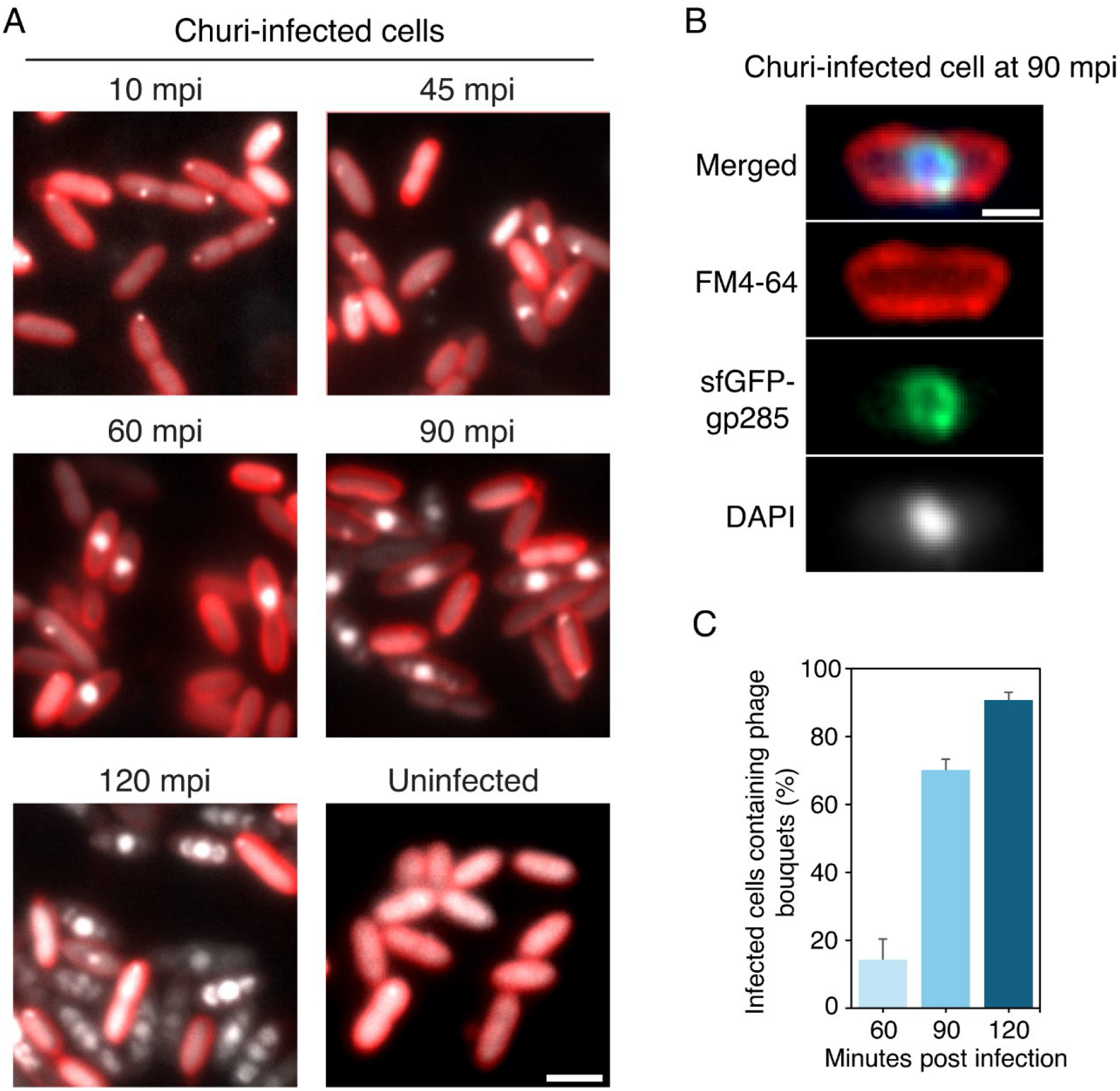
**(A)** Raw images of Churi infection (10, 45, 60, 90, and 120 mpi). FM4-64 (red) represents bacterial cell membrane and DAPI (gray) represent DNA. Scale bar represents 2 µm. **(B)** sfGFP-gp285 (ChmA) morphology at 90 mpi of Churi infection against *P. aeruginosa*. Scale bar is 1 µm. **(C)** Bouquet counts of phage Churi during infection (60, 90, and 120 mpi) against *P. aeruginosa* K2733. n≥150 cells in each mpi.

**Supplementary Figure 2.**
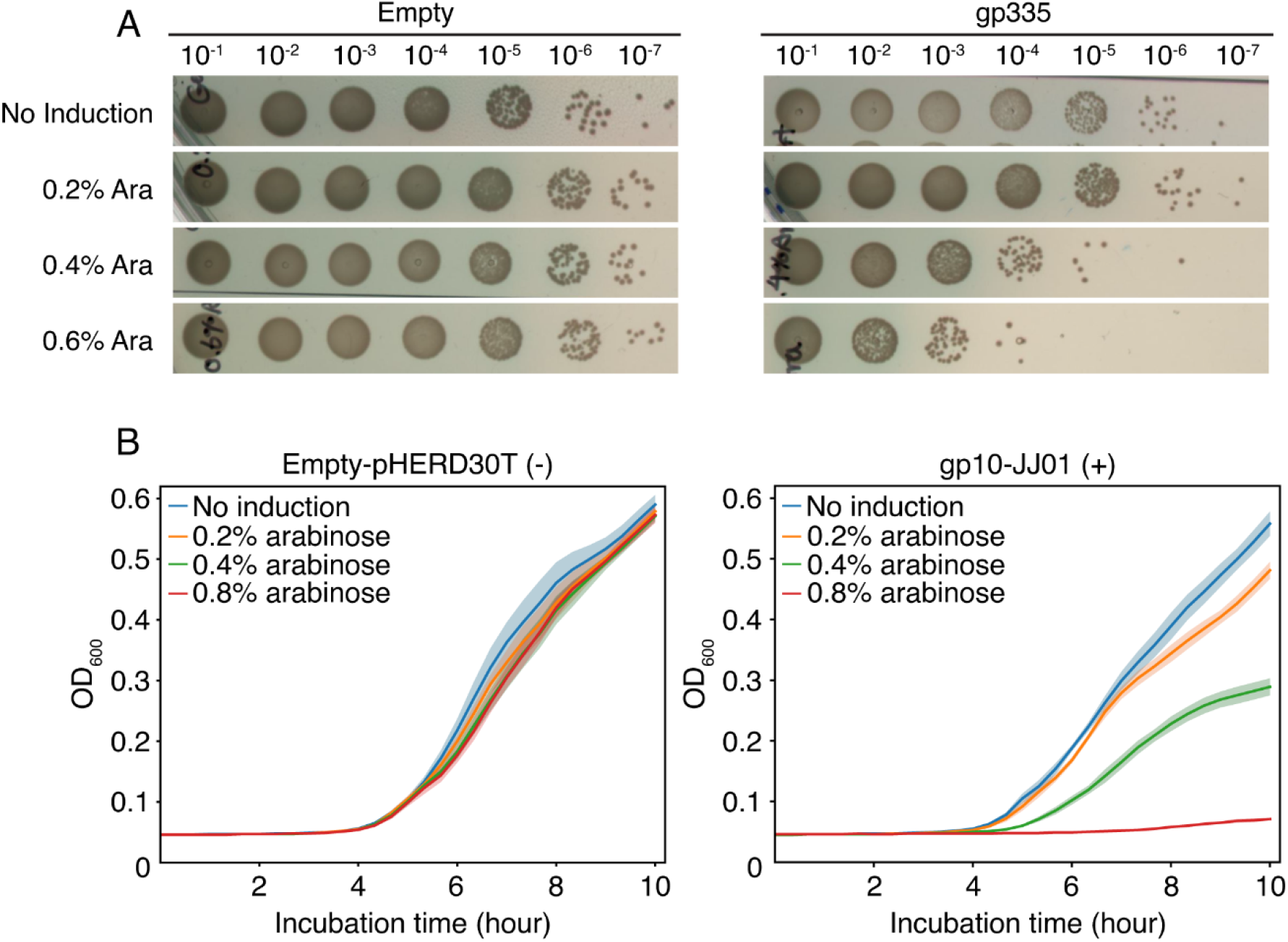
**(A)** Growth inhibition assay of gp335-Churi with different arabinose concentrations from 0.2 to 0.6% compared to empty-pHERD30T. **(B)** Optical density (OD600) of bacteria expressing empty-pHERD30T (negative control), and gp10-JJ01 (positive control) when the cells were induced with different concentration of arabinose (0.2, 0.4, and 0.8%). Shaded error bar represents standard deviation (±SD) of n=6.

**Supplementary Figure 3.**
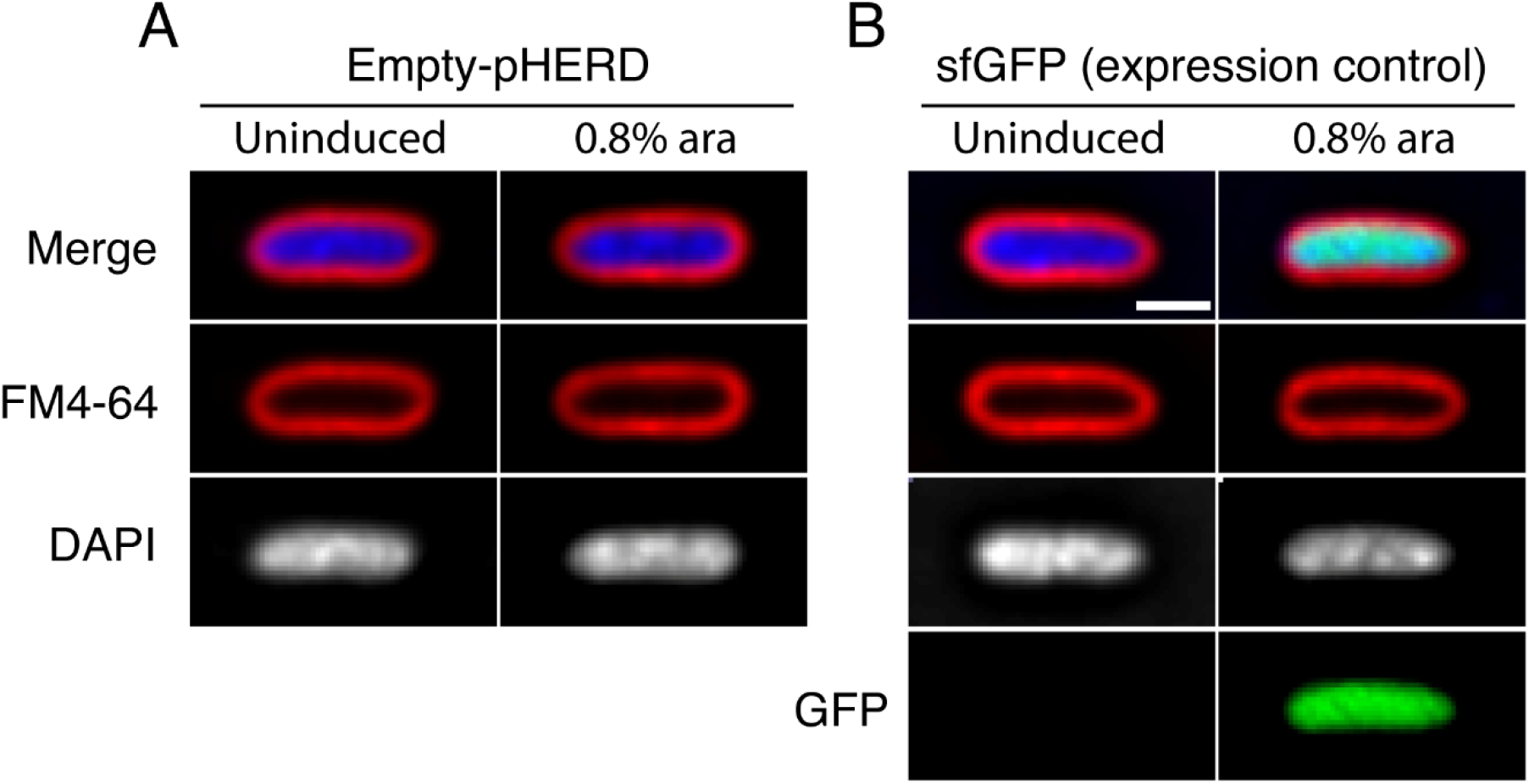
The fluorescence images of *P. aeruginosa* K2733 when induced with or without arabinose. **(A)** Empty-pHERD30T, and **(B)** sfGFP used as a gene expression control. Scale bar represents 1 µm.

**Supplementary Figure 4.**
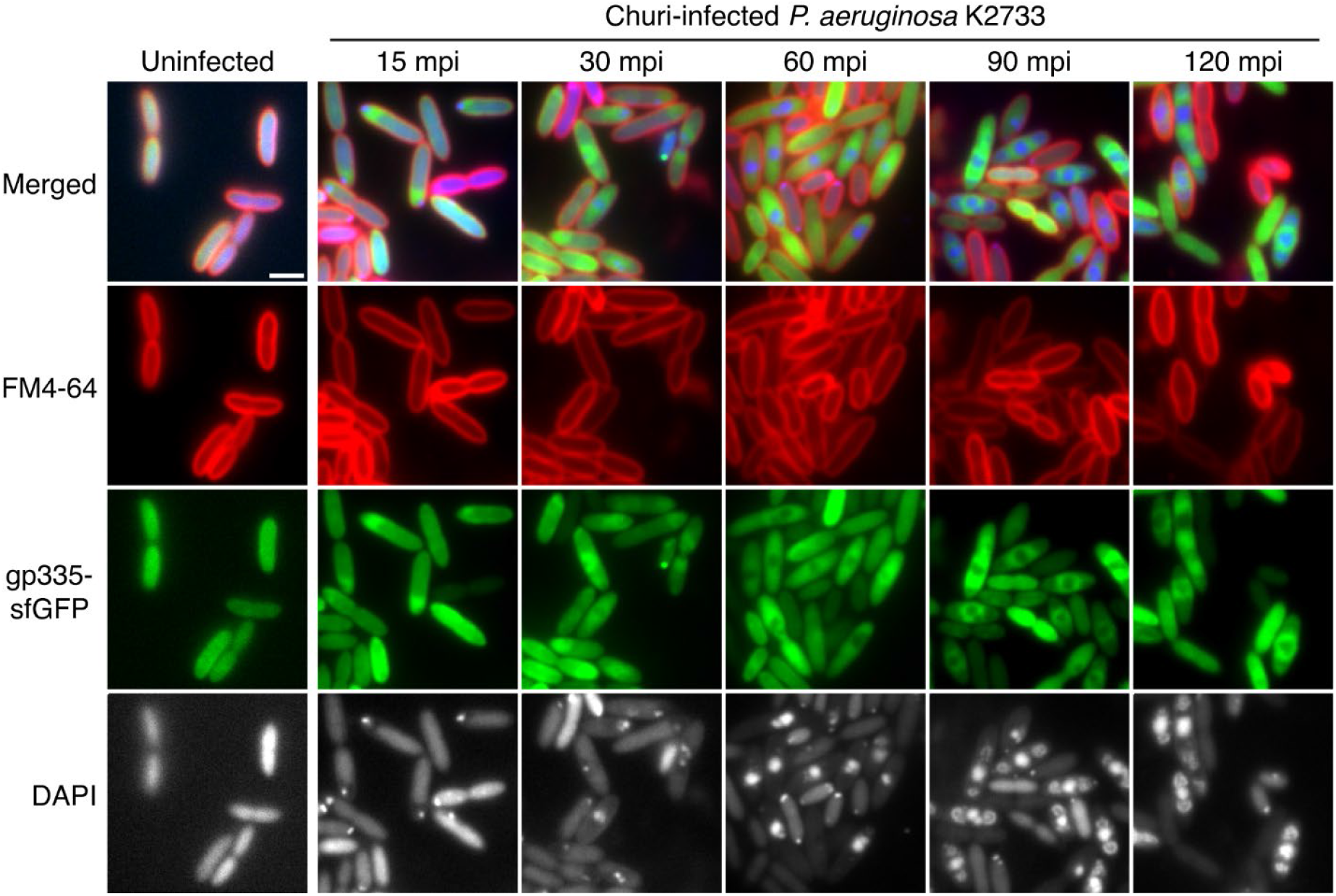
Raw images showing gp355-sfGFP localization when *P. aeruginosa* is infected with Churi (15 to 120 mpi). Scale bar represents 2 µm.

**Supplementary Figure 5.**
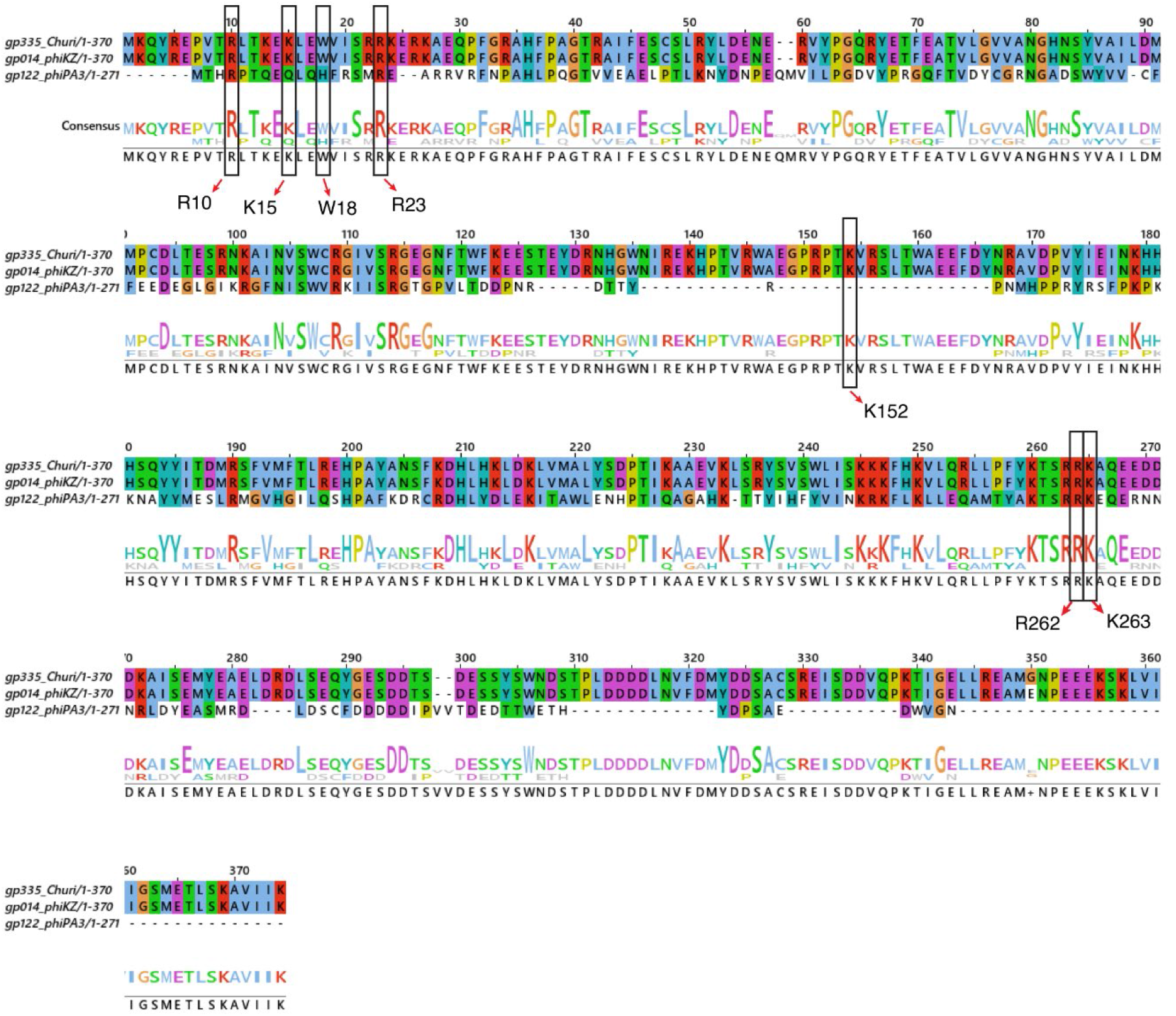
Multiple alignment of amino acid sequences (aligned with Clustal Omega analysis) between gp335-Churi against its homologs (gp014-phiKZ and gp122-phiPA3). Black frames represent the conserved amino acid residues that gp014-phiKZ uses to interact with host ribosomes as previously reported^12^.

**Supplementary Table 1.** VIRIDIC analysis of similarity distance (%) and cluster tables between Churi and other nucleus-forming phages (OMKO1, phiKZ, phiPA3, and 201phi2-1)

| **Genome** | Churi | OMKO1 | phiKZ | 201phi2-1 | PhiPA3 |
| --- | --- | --- | --- | --- | --- |
| Churi | 100 | 94.2 | 94.2 | 12.8 | 18.3 |
| OMKO1 | 94.2 | 100 | 94.4 | 12.5 | 18.2 |
| phiKZ | 94.2 | 94.4 | 100 | 12.3 | 18.5 |
| 201phi2-1 | 12.8 | 12.5 | 12.3 | 100 | 20.0 |
| PhiPA3 | 18.3 | 18.2 | 18.5 | 20.0 | 100 |

| **Genome** | **Species cluster** | **Genus cluster** |
| --- | --- | --- |
| Churi | 2 | 2 |
| OMKO1 | 3 | 2 |
| phiKZ | 4 | 2 |
| 201phi2-1 | 1 | 1 |
| PhiPA3 | 5 | 3 |

**Supplementary Table 2.** Mass Spectrometry results of Churi proteins that were detected during early infection against *P. aeruginosa* (**infection on plate**). Abundance indicates the number of peptides counted.

| Proteins | Function | Sum PEP Score | Coverage (%) | # Peptides | # PSMs | # Unique Peptides | Abundances (15 mpi) |
| --- | --- | --- | --- | --- | --- | --- | --- |
| gp005 | Hypothetical protein | 2.945 | 40 | 2 | 3 | 2 | 162.4 |
| gp039 | Hypothetical protein | 1.171 | 6 | 1 | 15 | 1 | 54.2 |
| gp055 | Hypothetical protein | 3.035 | 27 | 1 | 5 | 1 | 174 |
| gp059 | Hypothetical protein | 2.523 | 6 | 1 | 26 | 1 | 98.6 |
| gp071 | Hypothetical protein | 1.294 | 7 | 1 | 9 | 1 | 136 |
| gp094 | Hypothetical protein | 1.425 | 2 | 1 | 22 | 1 | 48.4 |
| gp110 | Hypothetical protein | 1.171 | 3 | 1 | 23 | 1 | 125.4 |
| gp119 | Hypothetical protein | 1.537 | 4 | 1 | 37 | 1 | 82.5 |
| gp123 | Hypothetical protein | 1.162 | 10 | 1 | 5 | 1 | 127.3 |
| gp126 | Hypothetical protein | 2.026 | 10 | 1 | 1 | 1 | 51 |
| gp130 | Hypothetical protein | 4.557 | 8 | 1 | 14 | 1 | 64.7 |
| gp135 | Hypothetical protein | 6.008 | 19 | 1 | 5 | 1 | 139.7 |
| gp136 | Hypothetical protein | 1.348 | 1 | 1 | 37 | 1 | 88.3 |
| gp147 | Hypothetical protein | 1.681 | 3 | 1 | 36 | 1 | 73.9 |
| gp150 | Hypothetical protein | 2.705 | 13 | 2 | 52 | 2 | 105 |
| gp177 | Hypothetical protein | 5.032 | 4 | 2 | 17 | 2 | 105.1 |
| gp199 | Hypothetical protein | 7.569 | 4 | 2 | 4 | 2 | 66.7 |
| gp202 | Hypothetical protein | 1.168 | 5 | 1 | 9 | 1 | 178.8 |
| gp218 | Hypothetical protein | 1.073 | 7 | 1 | 13 | 1 | 98.5 |
| gp234 | Hypothetical protein | 1.52 | 1 | 1 | 4 | 1 | 86 |
| gp256 | Hypothetical protein | 4.921 | 8 | 2 | 2 | 2 | 21.3 |
| gp279 | Hypothetical protein | 2.388 | 33 | 2 | 3 | 2 | 215.4 |
| gp287 | Hypothetical protein | 2.811 | 2 | 1 | 16 | 1 | 95.4 |
| gp291 | Hypothetical protein | 1.09 | 16 | 1 | 1 | 1 | 36.6 |
| gp325 | Hypothetical protein | 2.784 | 20 | 1 | 3 | 1 | 72.9 |
| gp335 | Hypothetical protein | 2.36 | 2 | 1 | 18 | 1 | 89.4 |
| gp354 | Hypothetical protein | 1.961 | 28 | 1 | 3 | 1 | 158.2 |

**Supplementary Table 3.**
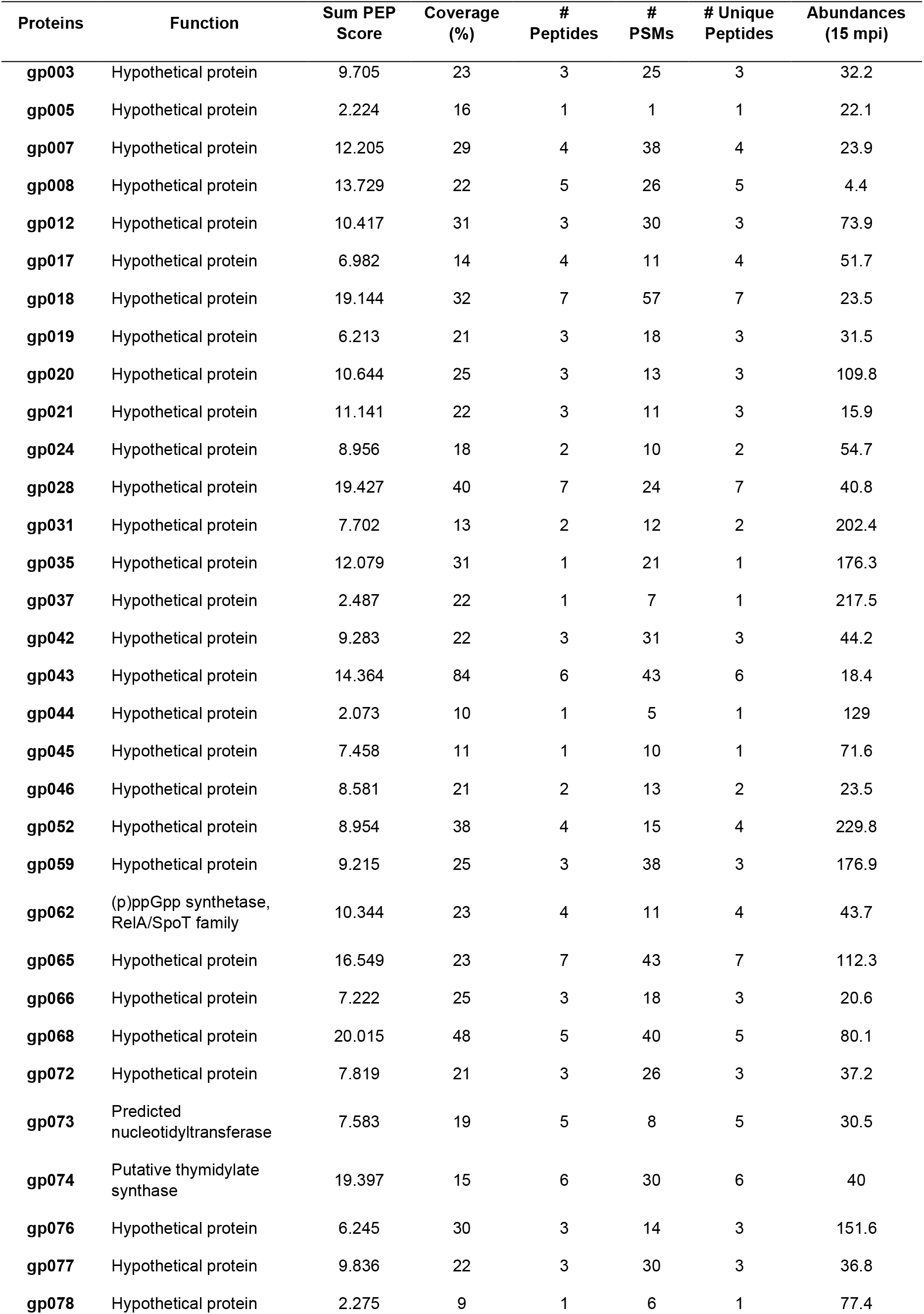
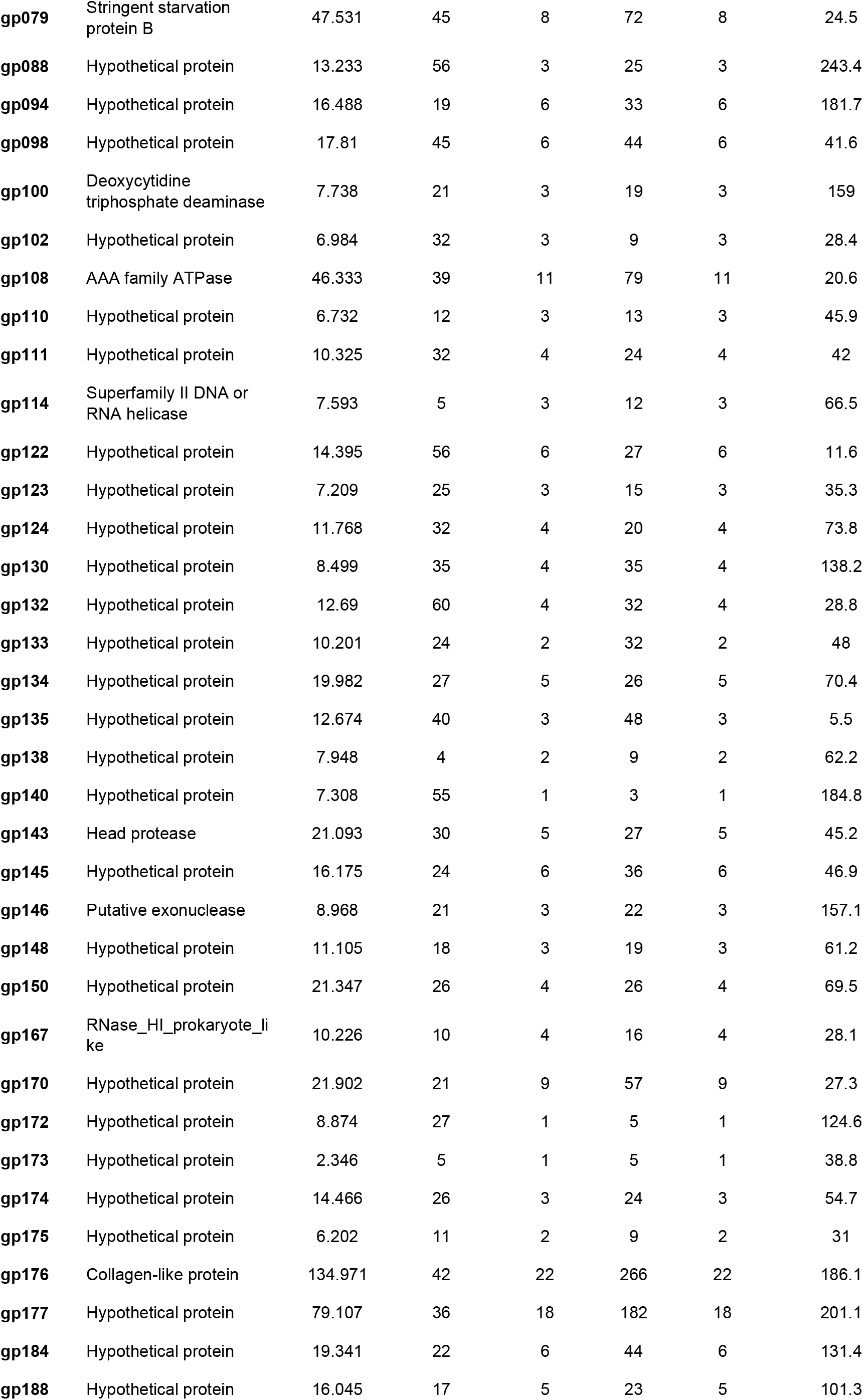
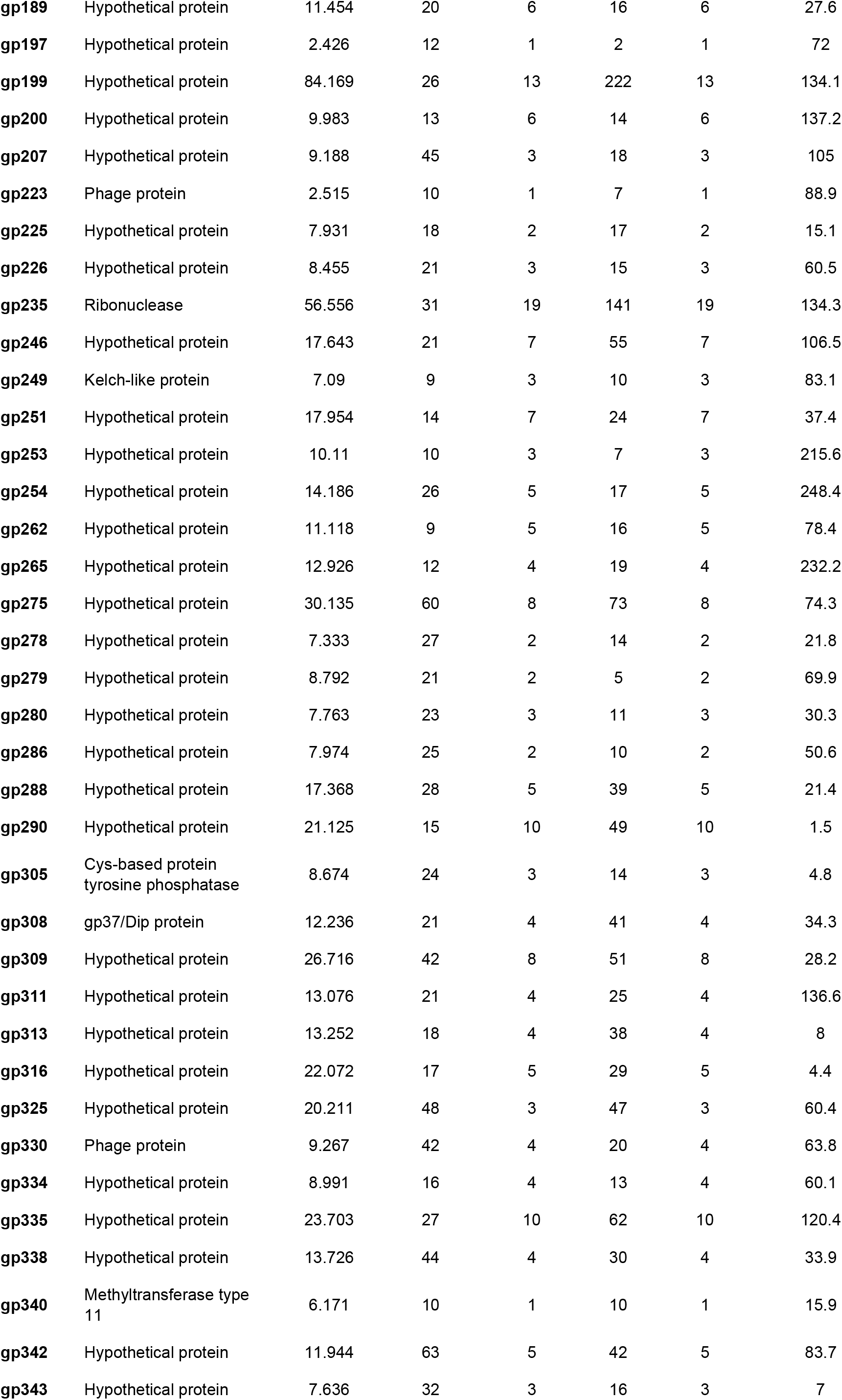
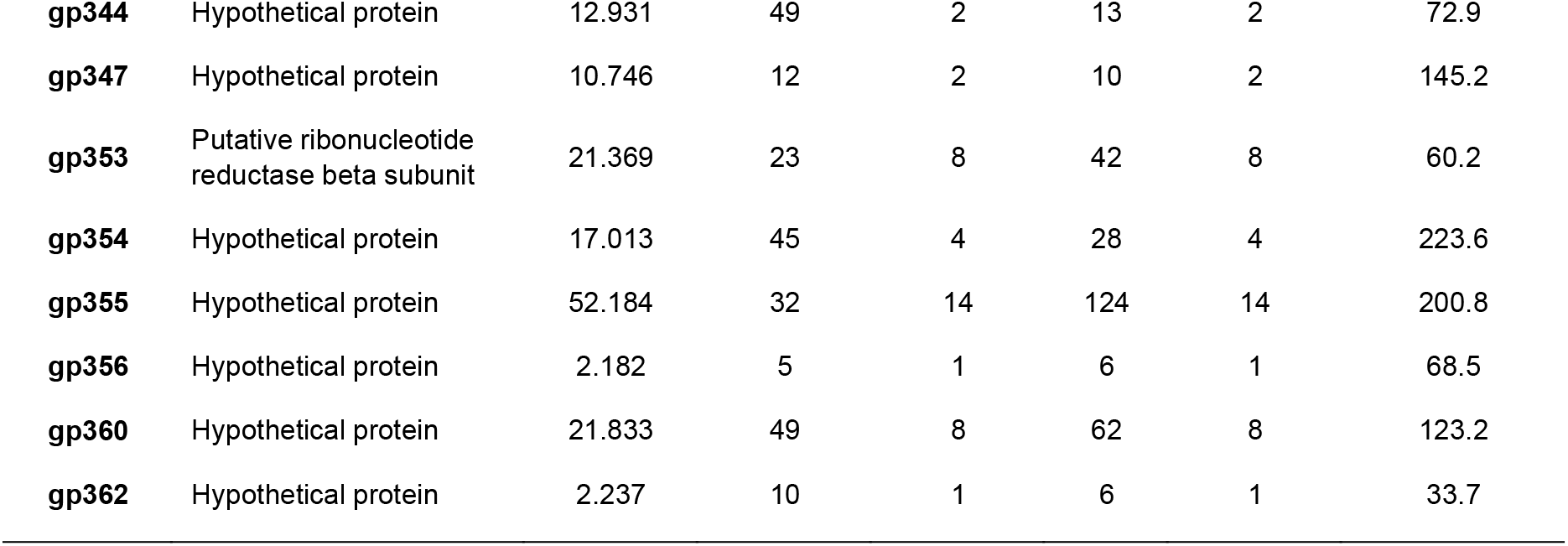
Mass Spectrometry results of Churi proteins that were detected during early infection against *P. aeruginosa* (**infection in broth**). Abundance indicates the number of peptides counted.

| Proteins | Function | Sum PEP Score | Coverage (%) | # Peptides | # PSMs | # Unique Peptides | Abundances (15 mpi) |
| --- | --- | --- | --- | --- | --- | --- | --- |
| gp003 | Hypothetical protein | 9.705 | 23 | 3 | 25 | 3 | 32.2 |
| gp005 | Hypothetical protein | 2.224 | 16 | 1 | 1 | 1 | 22.1 |
| gp007 | Hypothetical protein | 12.205 | 29 | 4 | 38 | 4 | 23.9 |
| gp008 | Hypothetical protein | 13.729 | 22 | 5 | 26 | 5 | 4.4 |
| gp012 | Hypothetical protein | 10.417 | 31 | 3 | 30 | 3 | 73.9 |
| gp017 | Hypothetical protein | 6.982 | 14 | 4 | 11 | 4 | 51.7 |
| gp018 | Hypothetical protein | 19.144 | 32 | 7 | 57 | 7 | 23.5 |
| gp019 | Hypothetical protein | 6.213 | 21 | 3 | 18 | 3 | 31.5 |
| gp020 | Hypothetical protein | 10.644 | 25 | 3 | 13 | 3 | 109.8 |
| gp021 | Hypothetical protein | 11.141 | 22 | 3 | 11 | 3 | 15.9 |
| gp024 | Hypothetical protein | 8.956 | 18 | 2 | 10 | 2 | 54.7 |
| gp028 | Hypothetical protein | 19.427 | 40 | 7 | 24 | 7 | 40.8 |
| gp031 | Hypothetical protein | 7.702 | 13 | 2 | 12 | 2 | 202.4 |
| gp035 | Hypothetical protein | 12.079 | 31 | 1 | 21 | 1 | 176.3 |
| gp037 | Hypothetical protein | 2.487 | 22 | 1 | 7 | 1 | 217.5 |
| gp042 | Hypothetical protein | 9.283 | 22 | 3 | 31 | 3 | 44.2 |
| gp043 | Hypothetical protein | 14.364 | 84 | 6 | 43 | 6 | 18.4 |
| gp044 | Hypothetical protein | 2.073 | 10 | 1 | 5 | 1 | 129 |
| gp045 | Hypothetical protein | 7.458 | 11 | 1 | 10 | 1 | 71.6 |
| gp046 | Hypothetical protein | 8.581 | 21 | 2 | 13 | 2 | 23.5 |
| gp052 | Hypothetical protein | 8.954 | 38 | 4 | 15 | 4 | 229.8 |
| gp059 | Hypothetical protein | 9.215 | 25 | 3 | 38 | 3 | 176.9 |
| gp062 | (p)ppGpp synthetase, RelA/SpoT family | 10.344 | 23 | 4 | 11 | 4 | 43.7 |
| gp065 | Hypothetical protein | 16.549 | 23 | 7 | 43 | 7 | 112.3 |
| gp066 | Hypothetical protein | 7.222 | 25 | 3 | 18 | 3 | 20.6 |
| gp068 | Hypothetical protein | 20.015 | 48 | 5 | 40 | 5 | 80.1 |
| gp072 | Hypothetical protein | 7.819 | 21 | 3 | 26 | 3 | 37.2 |
| gp073 | Predicted nucleotidyltransferase | 7.583 | 19 | 5 | 8 | 5 | 30.5 |
| gp074 | Putative thymidylate synthase | 19.397 | 15 | 6 | 30 | 6 | 40 |
| gp076 | Hypothetical protein | 6.245 | 30 | 3 | 14 | 3 | 151.6 |
| gp077 | Hypothetical protein | 9.836 | 22 | 3 | 30 | 3 | 36.8 |
| gp078 | Hypothetical protein | 2.275 | 9 | 1 | 6 | 1 | 77.4 |
| **gp079** | Stringent starvation protein B | 47.531 | 45 | 8 | 72 | 8 | 24.5 |
| **gp088** | Hypothetical protein | 13.233 | 56 | 3 | 25 | 3 | 243.4 |
| **gp094** | Hypothetical protein | 16.488 | 19 | 6 | 33 | 6 | 181.7 |
| **gp098** | Hypothetical protein | 17.81 | 45 | 6 | 44 | 6 | 41.6 |
| **gp100** | Deoxycytidine triphosphate deaminase | 7.738 | 21 | 3 | 19 | 3 | 159 |
| **gp102** | Hypothetical protein | 6.984 | 32 | 3 | 9 | 3 | 28.4 |
| **gp108** | AAA family ATPase | 46.333 | 39 | 11 | 79 | 11 | 20.6 |
| **gp110** | Hypothetical protein | 6.732 | 12 | 3 | 13 | 3 | 45.9 |
| **gp111** | Hypothetical protein | 10.325 | 32 | 4 | 24 | 4 | 42 |
| **gp114** | Superfamily II DNA or RNA helicase | 7.593 | 5 | 3 | 12 | 3 | 66.5 |
| **gp122** | Hypothetical protein | 14.395 | 56 | 6 | 27 | 6 | 11.6 |
| **gp123** | Hypothetical protein | 7.209 | 25 | 3 | 15 | 3 | 35.3 |
| **gp124** | Hypothetical protein | 11.768 | 32 | 4 | 20 | 4 | 73.8 |
| **gp130** | Hypothetical protein | 8.499 | 35 | 4 | 35 | 4 | 138.2 |
| **gp132** | Hypothetical protein | 12.69 | 60 | 4 | 32 | 4 | 28.8 |
| **gp133** | Hypothetical protein | 10.201 | 24 | 2 | 32 | 2 | 48 |
| **gp134** | Hypothetical protein | 19.982 | 27 | 5 | 26 | 5 | 70.4 |
| **gp135** | Hypothetical protein | 12.674 | 40 | 3 | 48 | 3 | 5.5 |
| **gp138** | Hypothetical protein | 7.948 | 4 | 2 | 9 | 2 | 62.2 |
| **gp140** | Hypothetical protein | 7.308 | 55 | 1 | 3 | 1 | 184.8 |
| **gp143** | Head protease | 21.093 | 30 | 5 | 27 | 5 | 45.2 |
| **gp145** | Hypothetical protein | 16.175 | 24 | 6 | 36 | 6 | 46.9 |
| **gp146** | Putative exonuclease | 8.968 | 21 | 3 | 22 | 3 | 157.1 |
| **gp148** | Hypothetical protein | 11.105 | 18 | 3 | 19 | 3 | 61.2 |
| **gp150** | Hypothetical protein | 21.347 | 26 | 4 | 26 | 4 | 69.5 |
| **gp167** | RNase_HI_prokaryote_like | 10.226 | 10 | 4 | 16 | 4 | 28.1 |
| **gp170** | Hypothetical protein | 21.902 | 21 | 9 | 57 | 9 | 27.3 |
| **gp172** | Hypothetical protein | 8.874 | 27 | 1 | 5 | 1 | 124.6 |
| **gp173** | Hypothetical protein | 2.346 | 5 | 1 | 5 | 1 | 38.8 |
| **gp174** | Hypothetical protein | 14.466 | 26 | 3 | 24 | 3 | 54.7 |
| **gp175** | Hypothetical protein | 6.202 | 11 | 2 | 9 | 2 | 31 |
| **gp176** | Collagen-like protein | 134.971 | 42 | 22 | 266 | 22 | 186.1 |
| **gp177** | Hypothetical protein | 79.107 | 36 | 18 | 182 | 18 | 201.1 |
| **gp184** | Hypothetical protein | 19.341 | 22 | 6 | 44 | 6 | 131.4 |
| **gp188** | Hypothetical protein | 16.045 | 17 | 5 | 23 | 5 | 101.3 |
| **gp189** | Hypothetical protein | 11.454 | 20 | 6 | 16 | 6 | 27.6 |
| **gp197** | Hypothetical protein | 2.426 | 12 | 1 | 2 | 1 | 72 |
| **gp199** | Hypothetical protein | 84.169 | 26 | 13 | 222 | 13 | 134.1 |
| **gp200** | Hypothetical protein | 9.983 | 13 | 6 | 14 | 6 | 137.2 |
| **gp207** | Hypothetical protein | 9.188 | 45 | 3 | 18 | 3 | 105 |
| **gp223** | Phage protein | 2.515 | 10 | 1 | 7 | 1 | 88.9 |
| **gp225** | Hypothetical protein | 7.931 | 18 | 2 | 17 | 2 | 15.1 |
| **gp226** | Hypothetical protein | 8.455 | 21 | 3 | 15 | 3 | 60.5 |
| **gp235** | Ribonuclease | 56.556 | 31 | 19 | 141 | 19 | 134.3 |
| **gp246** | Hypothetical protein | 17.643 | 21 | 7 | 55 | 7 | 106.5 |
| **gp249** | Kelch-like protein | 7.09 | 9 | 3 | 10 | 3 | 83.1 |
| **gp251** | Hypothetical protein | 17.954 | 14 | 7 | 24 | 7 | 37.4 |
| **gp253** | Hypothetical protein | 10.11 | 10 | 3 | 7 | 3 | 215.6 |
| **gp254** | Hypothetical protein | 14.186 | 26 | 5 | 17 | 5 | 248.4 |
| **gp262** | Hypothetical protein | 11.118 | 9 | 5 | 16 | 5 | 78.4 |
| **gp265** | Hypothetical protein | 12.926 | 12 | 4 | 19 | 4 | 232.2 |
| **gp275** | Hypothetical protein | 30.135 | 60 | 8 | 73 | 8 | 74.3 |
| **gp278** | Hypothetical protein | 7.333 | 27 | 2 | 14 | 2 | 21.8 |
| **gp279** | Hypothetical protein | 8.792 | 21 | 2 | 5 | 2 | 69.9 |
| **gp280** | Hypothetical protein | 7.763 | 23 | 3 | 11 | 3 | 30.3 |
| **gp286** | Hypothetical protein | 7.974 | 25 | 2 | 10 | 2 | 50.6 |
| **gp288** | Hypothetical protein | 17.368 | 28 | 5 | 39 | 5 | 21.4 |
| **gp290** | Hypothetical protein | 21.125 | 15 | 10 | 49 | 10 | 1.5 |
| **gp305** | Cys-based protein<br>tyrosine phosphatase | 8.674 | 24 | 3 | 14 | 3 | 4.8 |
| **gp308** | gp37/Dip protein | 12.236 | 21 | 4 | 41 | 4 | 34.3 |
| **gp309** | Hypothetical protein | 26.716 | 42 | 8 | 51 | 8 | 28.2 |
| **gp311** | Hypothetical protein | 13.076 | 21 | 4 | 25 | 4 | 136.6 |
| **gp313** | Hypothetical protein | 13.252 | 18 | 4 | 38 | 4 | 8 |
| **gp316** | Hypothetical protein | 22.072 | 17 | 5 | 29 | 5 | 4.4 |
| **gp325** | Hypothetical protein | 20.211 | 48 | 3 | 47 | 3 | 60.4 |
| **gp330** | Phage protein | 9.267 | 42 | 4 | 20 | 4 | 63.8 |
| **gp334** | Hypothetical protein | 8.991 | 16 | 4 | 13 | 4 | 60.1 |
| **gp335** | Hypothetical protein | 23.703 | 27 | 10 | 62 | 10 | 120.4 |
| **gp338** | Hypothetical protein | 13.726 | 44 | 4 | 30 | 4 | 33.9 |
| **gp340** | Methyltransferase type<br>11 | 6.171 | 10 | 1 | 10 | 1 | 15.9 |
| **gp342** | Hypothetical protein | 11.944 | 63 | 5 | 42 | 5 | 83.7 |
| **gp343** | Hypothetical protein | 7.636 | 32 | 3 | 16 | 3 | 7 |
| **gp344** | Hypothetical protein | 12.931 | 49 | 2 | 13 | 2 | 72.9 |
| **gp347** | Hypothetical protein | 10.746 | 12 | 2 | 10 | 2 | 145.2 |
| **gp353** | Putative ribonucleotide reductase beta subunit | 21.369 | 23 | 8 | 42 | 8 | 60.2 |
| **gp354** | Hypothetical protein | 17.013 | 45 | 4 | 28 | 4 | 223.6 |
| **gp355** | Hypothetical protein | 52.184 | 32 | 14 | 124 | 14 | 200.8 |
| **gp356** | Hypothetical protein | 2.182 | 5 | 1 | 6 | 1 | 68.5 |
| **gp360** | Hypothetical protein | 21.833 | 49 | 8 | 62 | 8 | 123.2 |
| **gp362** | Hypothetical protein | 2.237 | 10 | 1 | 6 | 1 | 33.7 |

**Supplementary Table 4.**
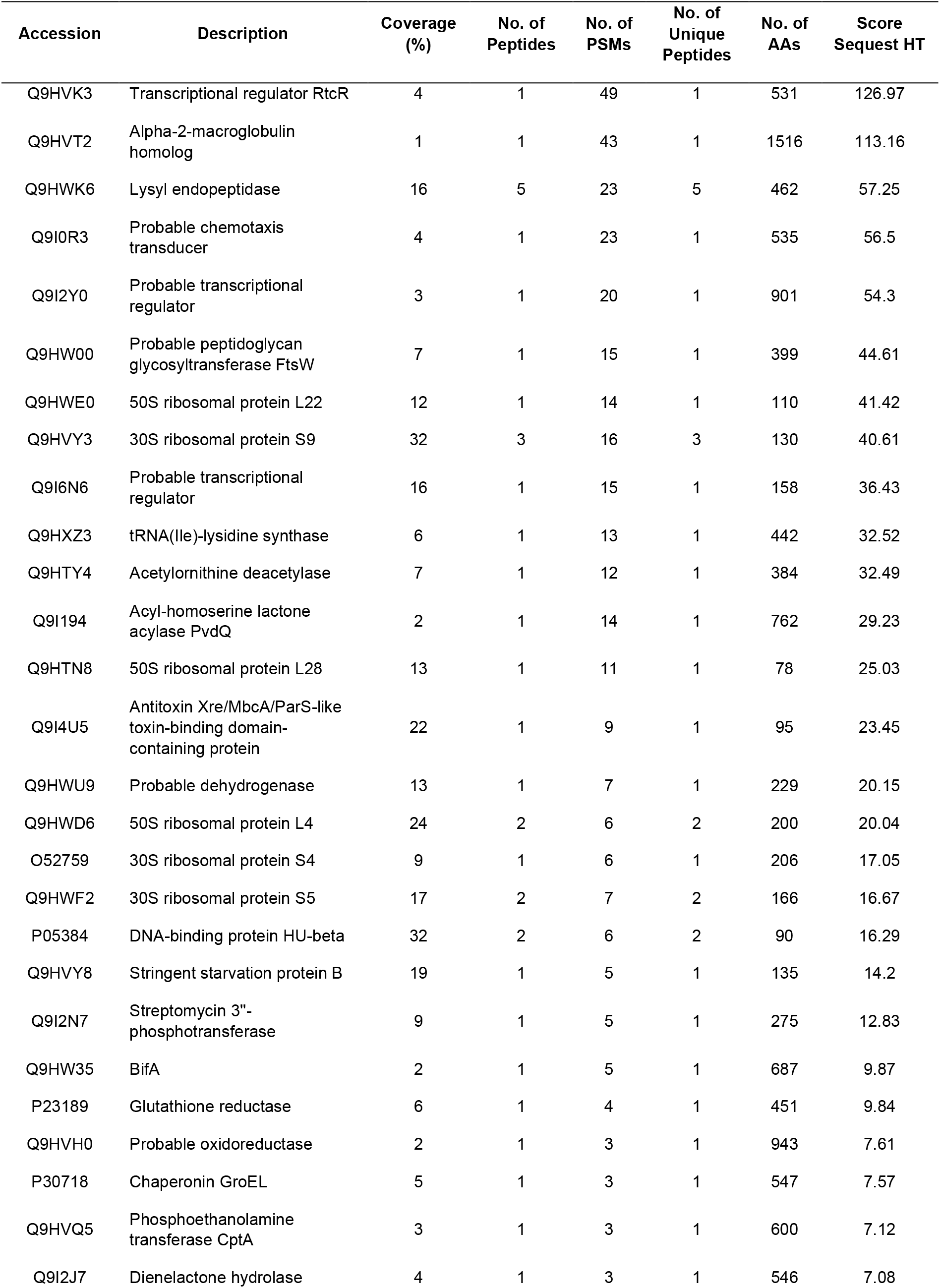
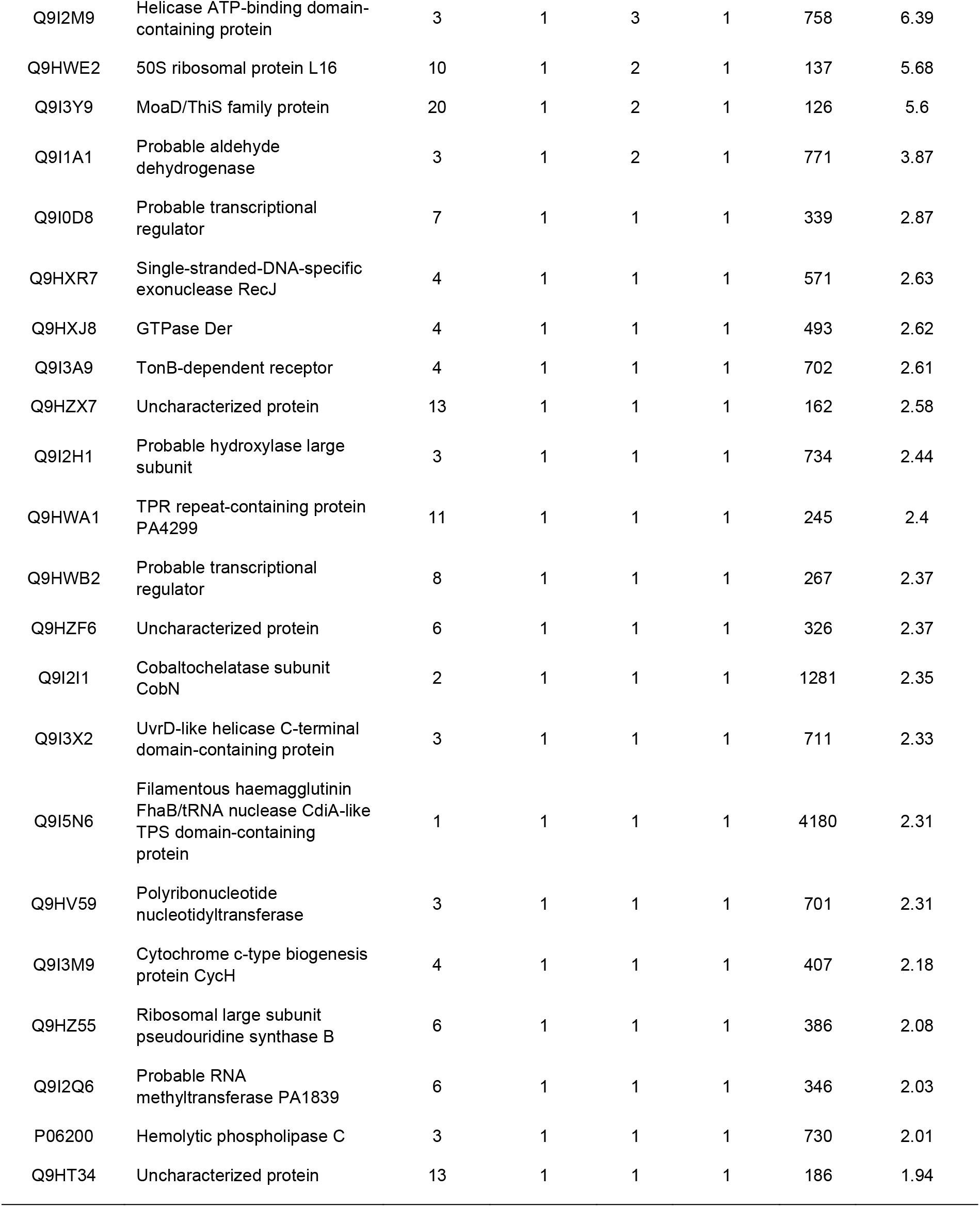
List of all proteins of *P. aeruginosa* PAO1 proteins according to Sequest HT score ranking that were detected from Co-immunoprecipitation experiment with gp335-sfGFP using Mass spectrometry.

**Supplementary Table 5.**
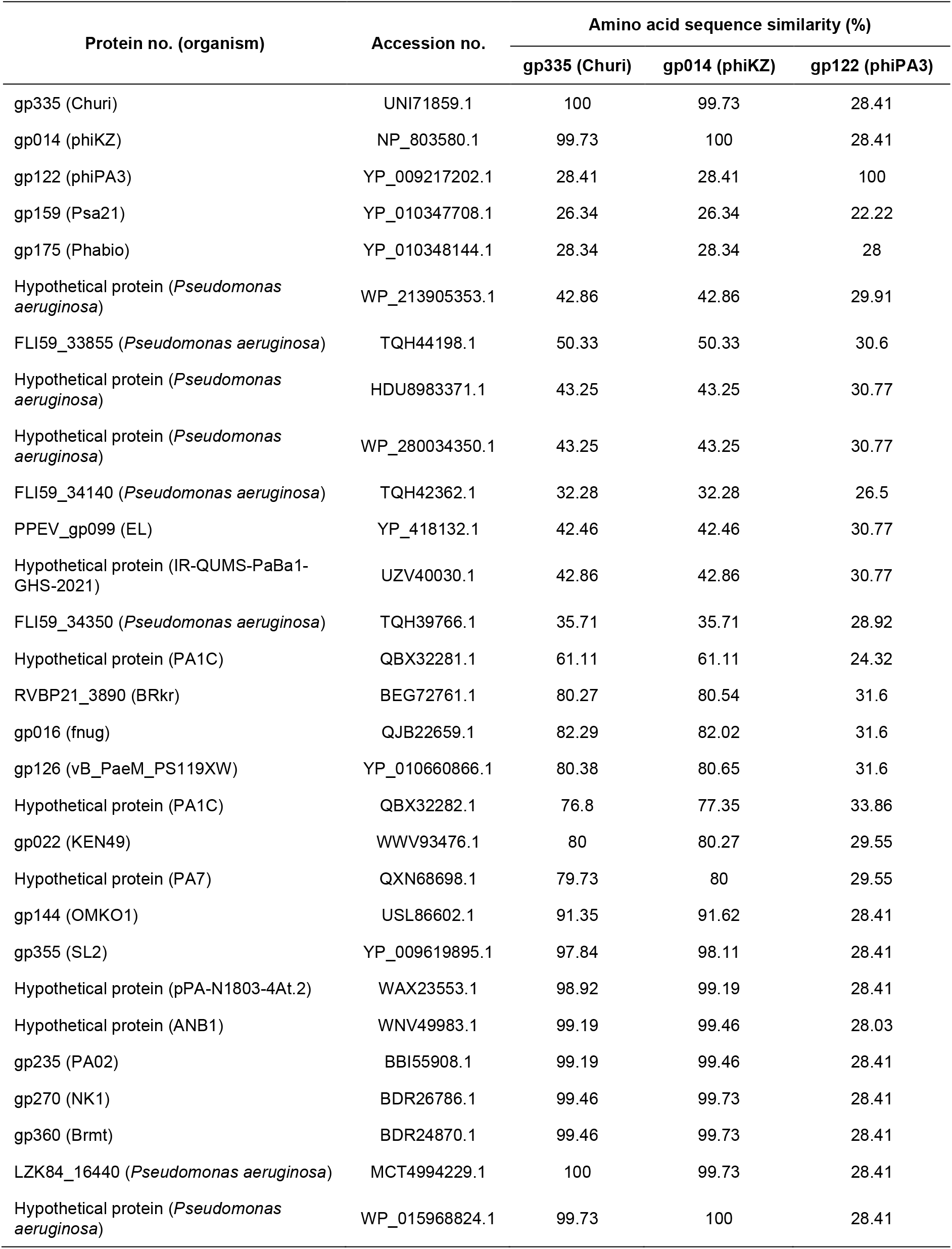
Amino acid sequence similarity (%) of gp335-Churi against its all homologs found in ncbi database (aligned with Clustal Omega analysis)

**Supplementary Table 6.** List of plasmid constructs that were used in this study.

| **Backbone vector** | **Insert** | **Strain No.** |
| --- | --- | --- |
| pHERD30T | sfGFP | WW1211 |
|  | JJ01-gp10 | WW1258 |
|  | Churi-gp005 | WW1080 |
|  | Churi-gp059 | WW1119 |
|  | Churi-gp094 | WW1178 |
|  | Churi-gp110 | WW1179 |
|  | Churi-gp123 | WW1181 |
|  | Churi-gp130 | WW1121 |
|  | Churi-gp135 | WW1085 |
|  | Churi-gp150 | WW1109 |
|  | Churi-gp177 | WW1183 |
|  | Churi-gp199 | WW1184 |
|  | Churi-gp256 | WW1186 |
|  | Churi-gp279 | WW1112 |
|  | Churi-gp325 | WW1149 |
|  | Churi-gp335 | WW1235 |
|  | Churi-gp354 | WW1087 |
|  | phiKZ-gp014 | WW1256 |
|  | phiPA3-gp122 | WW1257 |
|  | sfGFP-Churi-gp285 (ChmA) | WW1047 |
|  | Churi-gp335-sfGFP | WW1221 |
|  | Churi-gp335-guide-2 (CRISPR-Cas13a system) | WW1045 (Pogliano's lab) |
|  | Churi-gp335G2-2-sfGFP (mutant Churi-gp335) | WW1052 (Pogliano's lab) |

